# Coordinated repression of totipotency-associated gene loci by histone methyltransferase EHMT2 through binding to LINE-1 regulatory elements

**DOI:** 10.1101/2024.12.18.629181

**Authors:** K Chatterjee, CM Uyehara, K Kasliwal, S Madhuranath, L Scourzic, A Polyzos, E Apostolou, M Stadtfeld

**Affiliations:** Sanford I. Weill Department of Medicine, Sandra and Edward Meyer Cancer Center, Weill Cornell Medicine, New York, NY 10065, USA

**Author notes:** denotes equal contribution (K Chatterjee, CM Uyehara).

**Keywords:** 2-cell stage, H3K9 methylation, EHMT2, gene clusters, totipotency, pluripotency, retrotransposons, repressive domains

## Abstract

Mouse embryonic stem cells (mESCs) and other naïve pluripotent stem cells can reverse typical developmental trajectories and, at low frequency, de-differentiate into 2-cell-like cells (2CLCs) that resemble the mammalian embryo during zygotic genome activation (ZGA). This affords the opportunity to reveal molecular principles that govern the pre-implantation stages of mammalian development. We leveraged a multipurpose allele for acute protein depletion and efficient immunoprecipitation to dissect the molecular functions of the chromatin repressor EHMT2, a candidate antagonist of the mESC-to-2CLC transition. This allowed us to define categories of EHMT2 target genes characterized by distinct modes of EHMT2 chromatin engagement and repression. Most notably, EHMT2 directly represses large clusters of co-regulated gene loci that comprise a significant fraction of the 2CLC-specific transcriptome by initiating H3K9me2 spreading from distal LINE-1 elements. EHMT2 counteracts the recruitment of the activator DPPA2/4 to promoter-proximal endogenous retroviral elements (ERVs) at 2CLC genes. EHMT2 depletion elevates the expression of ZGA-associated transcripts in 2CLCs and synergizes with spliceosome inhibition and retinoic acid signaling in facilitating the mESC-to-2CLC transition. In contrast to ZGA-associated genes, repression of germ layer-associated transcripts by EHMT2 occurs outside of gene clusters in collaboration with ZFP462 and entails binding to non-repeat enhancers. Our observations show that EHMT2 attenuates the bidirectional differentiation potential of mouse pluripotent stem cells and define molecular modes for locus-specific transcriptional repression by this essential histone methyltransferase.

## INTRODUCTION

Mammalian stem and progenitor cells are unidirectional in their developmental plasticity and generally do not revert to earlier stages of differentiation. This is the basis for Waddington’s epigenetic landscape ^1^. Cultured naïve pluripotent stem cells, such as mouse embryonic stem cells (mESC), are characterized by their ability to differentiate into all three germ layers and derivative tissues. This reflects the developmental potential of their *in vivo* counterpart, the epiblast of the pre-implantation blastocyst. However, mESCs can also spontaneously reverse physiological developmental trajectories and, at low frequencies, give rise to so-called 2-cell-like cells (2CLCs)^2^. 2CLCs express genes that are usually only transiently activated during zygotic genome activation (ZGA), which in mice occurs at the two-cell stage before becoming silenced during later development ^2^. Several transcriptional regulators, such as the transcription factors DPPA2/4 ^3,4^ and DUX ^5,6^, have been reported to activate ZGA-associated genes during the mESC-to-2CLC transition.

The unusual bi-directional potential of mESCs suggests that molecular mechanisms exist that not only support but also counteract “forward” (into germ layers) and “backward” (into 2CLC) differentiation, thereby allowing mESCs to self-renew in an undifferentiated state. For example, core pluripotency transcription factors (TFs) such as OCT4 can recruit repressive chromatin modifiers to loci encoding signaling and transcriptional regulators required for germ layer differentiation ^7^. Several distinct cellular pathways and regulators have been reported to be involved in regulating the mESC-to-2CLC conversion ^8,9^, suggesting the existence of multiple regulatory layers that converge on suppressing the unscheduled re-activation of 2CLC-associated transcripts in pluripotent cells. However, the degree to which the same regulators are involved in counteracting either 2CLC formation or forward differentiation of mESCs remains unexplored.

The extensive differences in genome accessibility ^5^, histone mobility ^10^, chromatin marks ^11^, and chromatin topology ^12^ distinguishing mESCs and 2CLCs make epigenetic regulators prime candidates for modulating the interconversion between these cells. Accordingly, the inhibition of histone-modifying enzymes can increase the abundance of 2CLCs in mESCs cultures ^2^. However, the specific target genes of these enzymes and underlying regulatory mechanisms remain unknown.

Euchromatic histone methyltransferase 2 (EHMT2) was identified as the enzyme catalyzing the repressive H3K9me2 mark in gene-rich regions outside of the pericentromeric heterochromatin of the mammalian genome ^13,14^ together with its dimeric interaction partner EHMT1 ^15^. EHMT2 null mice die during early organogenesis with multi-lineage defects ^16^, but transcriptional dysregulation in the absence of EHMT2 is already evident at pre-implantation stages^17^. Since EHMT2 does not contain any DNA-binding domain, it is believed to gain target gene specificity by cell type-specific recruiting factors such as the TF ZFP462 in mESCs ^18^. Cultures of mESCs deficient for EHMT2 exhibit upregulation of gene loci associated with neurodevelopment and other germ layers ^19^, as well as ectopic activation of specific transposable elements (TE) highly expressed in two-cell embryos such as ERVs ^20^ and an elevated number of 2CLCs ^2^. Combined, these observations support a potential role of EHMT2 for counteracting both “forward” and “backward” differentiation in mESCs.

Here, we employ degron alleles to explore the molecular role of EHMT2 in mouse ESCs. Acute EHMT2 depletion reveals that a significant fraction (∼30%) of 2CLC-associated transcripts arise from co-regulated gene clusters that we term “EHMT2 coordinately repressed domains” (ECORDs). Genes within ECORDs are highly expressed during ZGA *in vivo*, and we show that loss of EHMT2 further elevates ZGA-associated transcripts in 2CLCs, resulting in the inability of these cells to revert to a pluripotent state. An antagonism between EHMT2 and the activating transcription factor DPPA2/4 regulates the stage-specific expression of ECORDs via recruitment to distinct types of transposable elements (TEs). At the same time, EHMT2 synergizes with ZFP462 to silence differentiation-associated genes. Our study defines distinct gene regulatory modes that EHMT2 engages in to preserve the remarkable developmental plasticity of naïve pluripotent stem cells.

## RESULTS

### Acute EHMT2 depletion de-represses distinct categories of gene loci in mESCs

To facilitate the study of EHMT2’s gene regulatory functions, we replaced its STOP codon in mouse embryonic stem cells (mESCs) with an in-frame transgenic cassette encoding the degron tag FKBP12^F36V^ (“dTAG”) ^21^ and two copies of the hemagglutinin (HA) tag (**Fig.1A**). The degron design also contains a mCherry reporter to capture changes in the transcriptional activity of *Ehmt2* and to facilitate isolation of correctly targeted cells (**Fig.1A**). We generated several PCR-validated homozygous EHMT2-dTAG mESC lines. Flow cytometric analysis of the mCherry reporter confirmed homogeneous *Ehmt2* expression in pluripotent cells (**Fig. S1A**). Culture of EHMT2-dTAG mESCs for 24 hours (h) in the presence of the degrader dTAG-13 ^21^ resulted in near complete elimination of EHMT2 protein as measured by Western Blot (**Fig. S1B**). Quantification by flow cytometry revealed that total EHMT2 depletion was achieved after 6h of dTAG-13 treatment, followed by a delayed reduction in the levels of the H3K9me2 mark, which is catalyzed by the EHMT1:EHMT2 complex (**Fig.1B** and **Fig.S1C**). No reduction of H3K9me2 levels was observed in EHMT2-dTAG mESCs in the absence of dTAG-13 (**Fig.S1D**). Thus, our transgenic system achieves robust, dTAG-13-dependent control of EHMT2 levels in mESCs.

**Figure 1.**
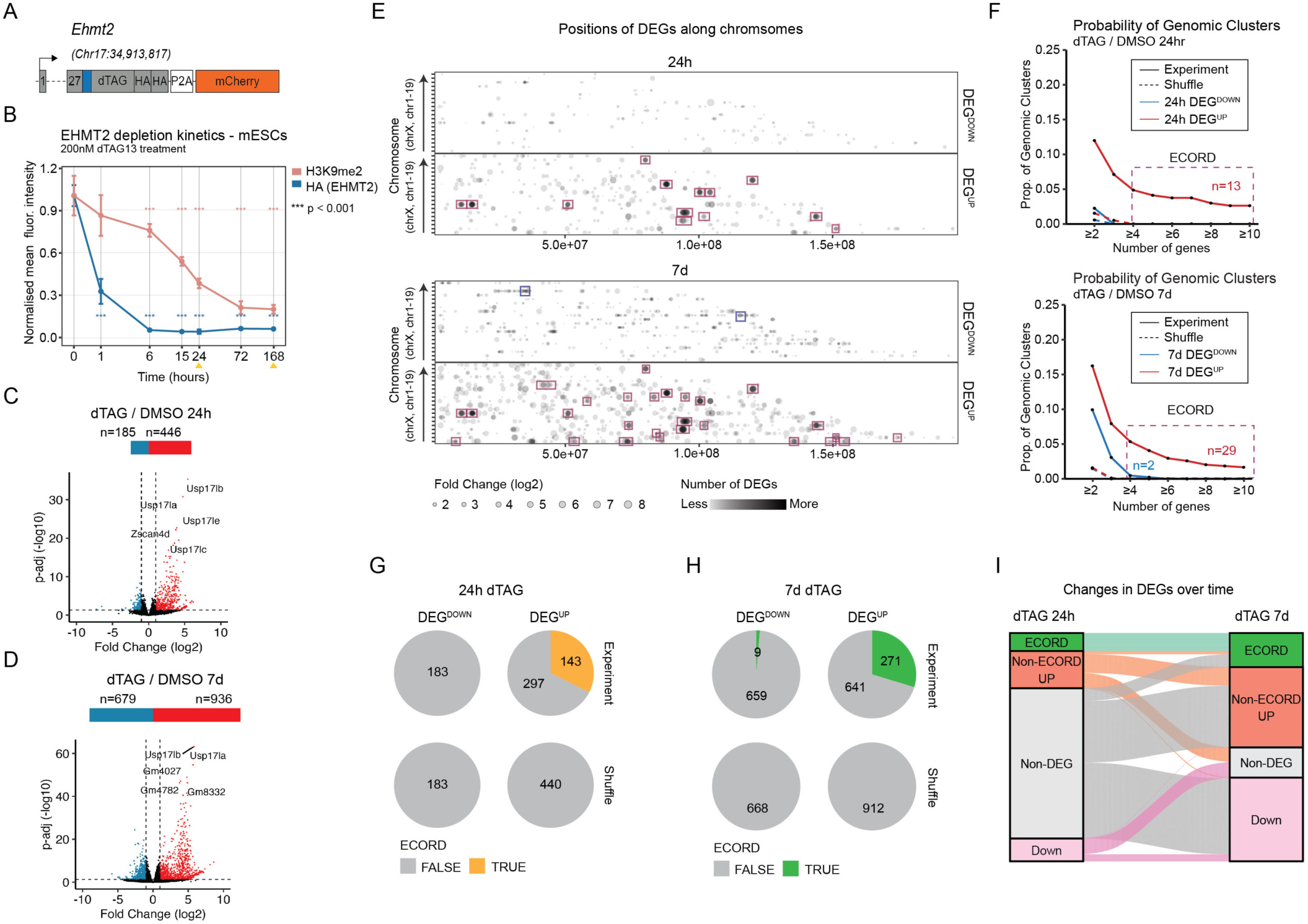
Acute EHMT2 depletion reveals distinct categories of target genes in mESCs. **A)** Schematic of the EHMT2 degron allele. A degron (dTAG), two HA tags, and a mCherry transcriptional reporter were integrated in-frame at the single STOP codon of the endogenous *Ehmt2* locus. **B)** Flow cytometry analysis of EHMT2 (HA) and H3K9me2 levels over time in response to continuous dTAG treatment. Yellow arrows indicate the isolation of samples for RNA-seq analysis. **C,D)** Volcano plots showing numbers of differentially expressed genes (DEGs) following 24h and 7d continuous dTAG treatment, respectively. DEG: Adj. p-value < 0.05 and absolute log2 fold change ≥1. The top 5 DEGs by fold change are highlighted. **E)** Positions of 24h and 7d EHMT2 DEGs along the linear genome. The opacity of the points was decreased so areas with large numbers of high FC DEGs appear darker. Boxes highlight genes (red: upregulated, blue: downregulated) that fall into EHMT2 Coordinately Repressed Domains (ECORDs) as defined in panels F and S1H-I. **F)** Probability curves showing the fraction of gene clusters with ≥n DEGs and < 50% static genes that do not break TAD boundaries (see Results and Methods sections). For subsequent analyses, “ECORDs” were defined as clusters with n ≥4 DEGs and <0.5 proportion of static genes. **G,H)** Fraction of DEGs that fall into ECORDs at 24h and 7d, respectively. In panels F-G, “Shuffle” refers to a random sample of genes expressed in either DMSO or dTAG conditions. **I)** Alluvial plot showing changes in DEGs between 24h and 7d.

To determine the transcriptional consequences of acute EHMT2 depletion, we conducted RNA-sequencing (RNA-seq) experiments with three independent EHMT2-dTAG cell clones 24h after dTAG-13 administration. This revealed 631 differentially expressed genes (DEGs) (abs(log2FC)>1; p-adj <0.05) compared to DMSO-treated controls (**Fig.1C**). Consistent with a predominant role of EHMT2 as a transcriptional repressor in mESCs ^22,23^, most DEGs (446/631 or 70.7%) were up-regulated and up-regulated DEGs also had higher fold changes than down-regulated DEGs (**Fig.1C**). To determine the longer-term consequences of EHMT degradation, we also conducted RNA-seq 7 days (d) after continuous dTAG-13 administration. Although mESC treated in this manner remained viable and retained an undifferentiated morphology, 7d RNA-seq revealed a substantially more pronounced transcriptional effect of EHMT2 loss with a total of 1,615 DEGs, a slight majority being upregulated (58.0% or 936/1,615 genes) (**Fig. 1D**) (**Table S1**). Previous work has reported that EHMT2 antagonizes the expression of specific endogenous retroviral transcripts ^20^. In line with this, EHMT2 depletion resulted in dysregulation of several repeat elements with upregulation of specific ERVK/ERVL LTR families among the earliest and most pronounced consequences (**Fig.S1E**). At 7d, we observed further de-repression of ERVs, which is in line with a significant role of EHMT2 in stably repressing these repeat families (**Fig.S1F**). Most DEGs upregulated (“DEG^UP^”) upon prolonged EHMT2 depletion already showed at least a trend towards upregulation after acute EHMT2 depletion (**Fig.S1G**). However, we also observed subsets of DEG^UP^ that were specific to the 24h (C5 in **Fig.S1G**) or 7d (C1 and C6) timepoint, possibly suggesting the existence of compensatory repressive mechanisms or the accumulation of indirect molecular effects of EHMT2 loss, respectively.

A striking outcome of our RNA-seq analysis was that a subset of DEG^UP^ was in linear proximity to one another and formed apparent clusters of genes that all showed a similar response to EHMT2 depletion. By projecting the chromosomal locations and fold-changes of 24h and 7d DEGs, we confirmed that many strongly upregulated (but not downregulated genes) were organized into clusters along the linear genome, suggesting coordinated repression by EHMT2 (**Fig.1E**). To identify clusters in an unbiased and quantitative fashion, we counted the number of DEGs that occurred in sequence along the linear genome. The cluster was interrupted if 1) the next DEG changed in the opposite direction (p-adj <0.05, no fold-cutoff), or 2) the cluster crossed a TAD boundary ^24^. This analysis revealed that DEG^UP^ clusters spanned across larger genomic regions than DEG^DOWN^ clusters and contained a much lower proportion of static (expressed and p-adj >0.05) genes (**Fig. 1F, S1H).** In some cases, >20 DEG^UP^ occurred in sequence with only 1-2 static genes. This analysis confirmed that clustering is a feature unique to a subset of DEG^UP^ that does not happen to DEG^DOWN^ or through chance. We will refer to genomic clusters with ≥4 DEG^UP^ and <50% static genes as **<u>E</u>**HMT2 **<u>Co</u>**ordinately **<u>R</u>**epressed **<u>D</u>**omains or <u>ECORD</u>s. Overall, we identified 13 ECORDs at 24h and 29 ECORDs at 7d (**Fig. 1F, S1J**), which made up 32.1% and 29.0% of DEG^UP^, respectively, at these two time points (**Fig.1G,H**) (**Table S2**). All 24h ECORDs were maintained at 7d (**Fig1I and Fig.S1K**), and the vast majority (24 out of 29) of 7d ECORDs had at least one DEG^UP^ at 24h, demonstrating that EHMT2 loss causes early and sustained upregulation of genes within ECORDs. In contrast, some 24h DEG^UP^ outside of ECORDs (“non-ECORD DEGs”) were no longer identified as DEG^UP^ at 7d, possibly suggesting the existence of compensatory repressive mechanisms at non-clustered gene loci (**Fig1I**).

### ECORDs are characterized by repressive chromatin and are associated with zygotic genome activation

The results described so far show that genes repressed by EHMT2 in mESC cultures can be separated into two broad groups based on whether they reside within or outside ECORDs. To elucidate potential regulatory differences between these groups, we analyzed their chromatin states with ChromHMM ^25,26^. ECORD genes strongly enriched for heterochromatin features such as H3K9me3, transposable elements (including LTRs and LINE-1 elements), and for the “Assembly Gap” class, which is characteristic of repetitive DNA (**Figs.2A and S2A; Table S3**).

In contrast, non-ECORD DEG^UP^ and DEG^DOWN^ enriched for chromatin features associated with euchromatin and active transcription (**Figs.2A and S2A; Table S3**). Accordingly, analysis of public Hi-C data ^24^ showed that ECORDs preferentially localized within (inactive) B compartments ^27^ while non-ECORD DEG^UP^ showed a weak enrichment for the A compartments at both 24h and 7d (**Fig.2B**). Collectively, these observations suggest that ECORD DEGs represent a more repressed ground state than non-ECORD DEG^UP^ in mESCs. In agreement, ECORD DEG^UP^ showed a more substantial degree of upregulation upon EHMT2 loss than non-ECORD DEG^UP^ (**Fig.2C**). Importantly, the expression of almost all ECORD DEGs—as well as non-ECORD DEGs—reverted to physiological levels upon dTAG-13 washout and EHMT2 recovery (**Fig.S2B-D**), demonstrating that EHMT2 is directly responsible for the repression of these loci and can regain transcriptional control after being transiently depleted.

**Figure 2.**
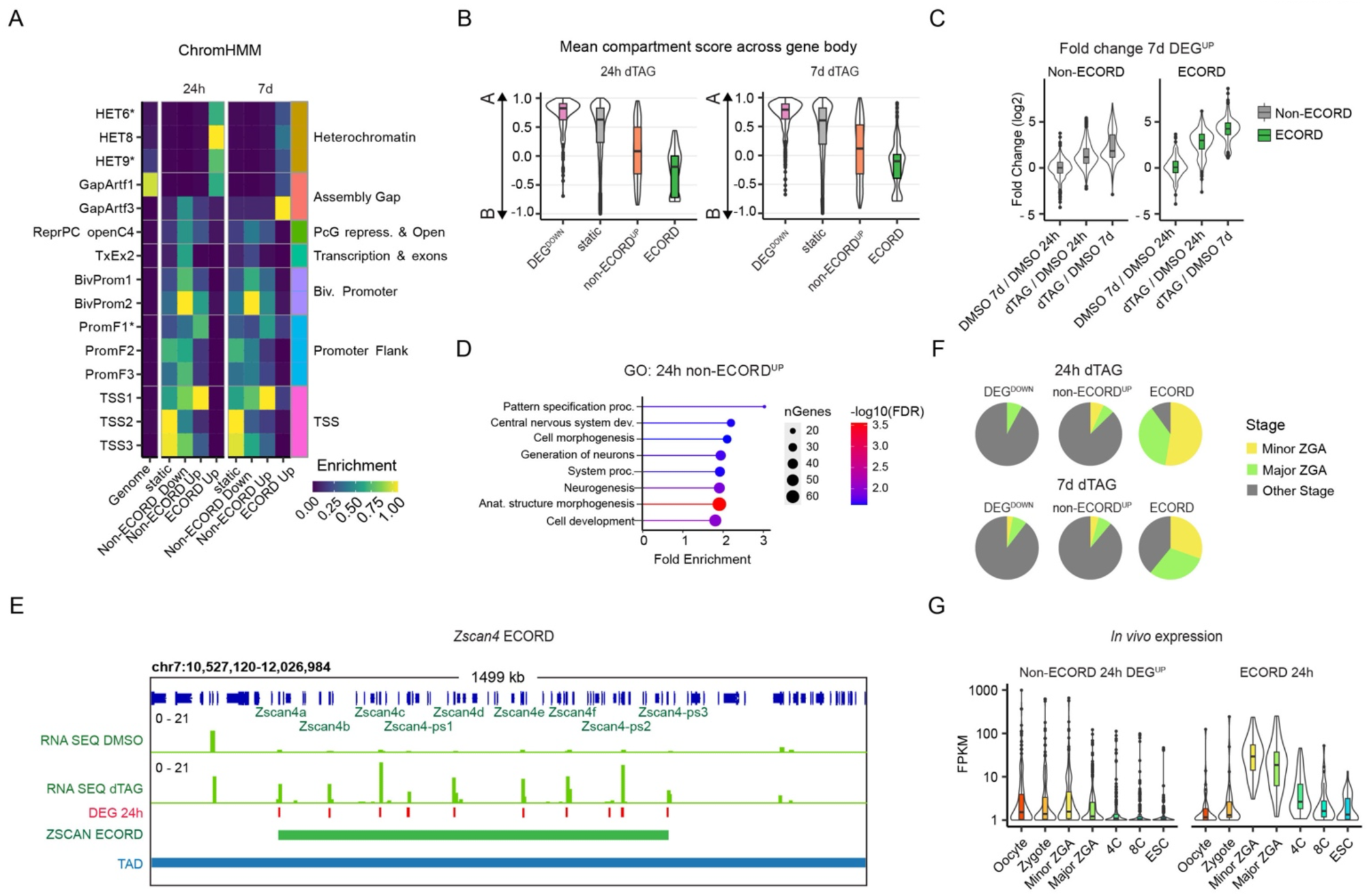
Genes in ECORDs have distinct properties and are enriched for genes involved in Zygotic Genome Activation (ZGA). **A)** ChromHMM of promoters of ECORD DEGs using a 100-state model ^26^. The descriptions of starred terms are provided in Fig.S2A. **B)** Mean compartment strength over different categories of genes following 24h and 7d dTAG treatment. Compartment scores were obtained from GSE113431 ^24^. **C)** Fold-changes (log2) of DEGs up-regulated after 7d dTAG treatment split by whether the DEG is within an ECORD. **D)** Gene Ontology (GO) terms of genes up-regulated after 24h dTAG treatment for Non-ECORD DEGs. **E)** Browser shot of RNA levels across the ECORD encompassing the *Zscan4* genes, canonical markers. TAD boundaries were obtained from GSE113339 ^24^. **F)** Overlap of 24h DEGs with gene categories from an *in vivo* time course of pre-implantation embryonic development ^30^. Only genes that overlapped annotated categories were included. **G)** Expression of 24h DEGs at the indicated stages of *in vivo* development and mESCs. Only genes with an FPKM ≥10 in at least one sample type were included.

To understand the potential biological relevance of genes repressed by EHMT2 in mESCs, we performed a Gene Ontology (GO) analysis. This showed that non-ECORD 24h DEG^UP^ were associated with developmental processes such as morphogenesis, neurogenesis, and organ development (**Fig.2D, Table S5**), which is consistent with the notion that EHMT2 functions in pluripotent cells to repress the premature expression of genes with regulatory roles during post-implantation stages of development. This agrees with the embryonic lethality of EHMT2 KO mice during organogenesis ^15,16^. In contrast, ECORD DEGs were not associated with post-implantation development but enriched for regulators of RNA localization and nuclear transport (**Fig.S2E, Table S5**). We noticed that ECORDs included genes known to become activated during zygotic genome activation (ZGA), such as the *Zscan4* ^28^ (**Fig.2E**) and *Obox* ^29^ loci. To further explore this, we compared our DEGs to a published dataset that characterized stage-specific transcripts during early mouse embryogenesis *in vivo* ^30^. Indeed, both 24h and 7d ECORD DEG^UP^ strongly overlapped with ZGA-associated transcripts and, in mouse embryos, are expressed at the highest levels during the early 2-cell (minor ZGA) stage, concomitant with the onset of ZGA (**Fig. 2F,G).** Non-ECORD DEG^UP^ did not exhibit this pattern (**Fig. 2G)**.

Together, these results support the notion that EHMT2 represses at least two broadly distinct categories of target gene loci in mouse pluripotent cells: clustered ECORD genes within heterochromatic regions which are transiently activated during ZGA and non-clustered, euchromatic loci encoding genes involved in later developmental stages.

### EHMT2 limits the entry rate of mESCs into a 2-cell-like transcriptional state in collaboration with other pathways

Cultures of mESCs can contain a small percentage of cells in a transient, 2-cell-like (2CLC) state characterized by the high-level expression of ZGA-associated genes such as *Zscan4* ^8,9^. Therefore, we hypothesized that the widespread up-regulation of ZGA genes in ECORDs we observed might reflect a change in the composition of our cell cultures to contain a higher percentage of 2CLCs. To enable dissecting the role of EHMT2 in controlling the emergence of 2CLCs and, ultimately, the control of ZGA-associated transcription, we generated EHMT2-dTAG mESC lines carrying destabilized, fast-folding TurboGFP reporters driven from murine endogenous retrovirus-L (MERVL) promoter elements ^31^ (“MERVL-GFP mESCs”) (**Fig.3A**). The activation of MERVL repeats is a hallmark of ZGA and is an established approach to identifying 2CLCs in culture ^8,9^. Furthermore, EHMT2 has been shown to repress MERVL elements in mESCs ^20^, a finding confirmed by our RNA-seq analysis of bulk cultures (see **Fig.S1E,F**). MERVL-GFP^+^ cells expressed the ZGA-associated, ECORD-encoded (**Fig.2E**) transcription factor (TF) ZSCAN4 (**Figs.3B, S3A**) and exhibited strongly reduced levels of the pluripotency-associated surface markers SSEA-1 and EpCAM ^32^ (**Fig.3C**). Hereafter, we refer to MERVL-GFP^+^ cells as “2CLCs” and to MERVL-GFP^-^ cells as “mESCs”.

**Figure 3.**
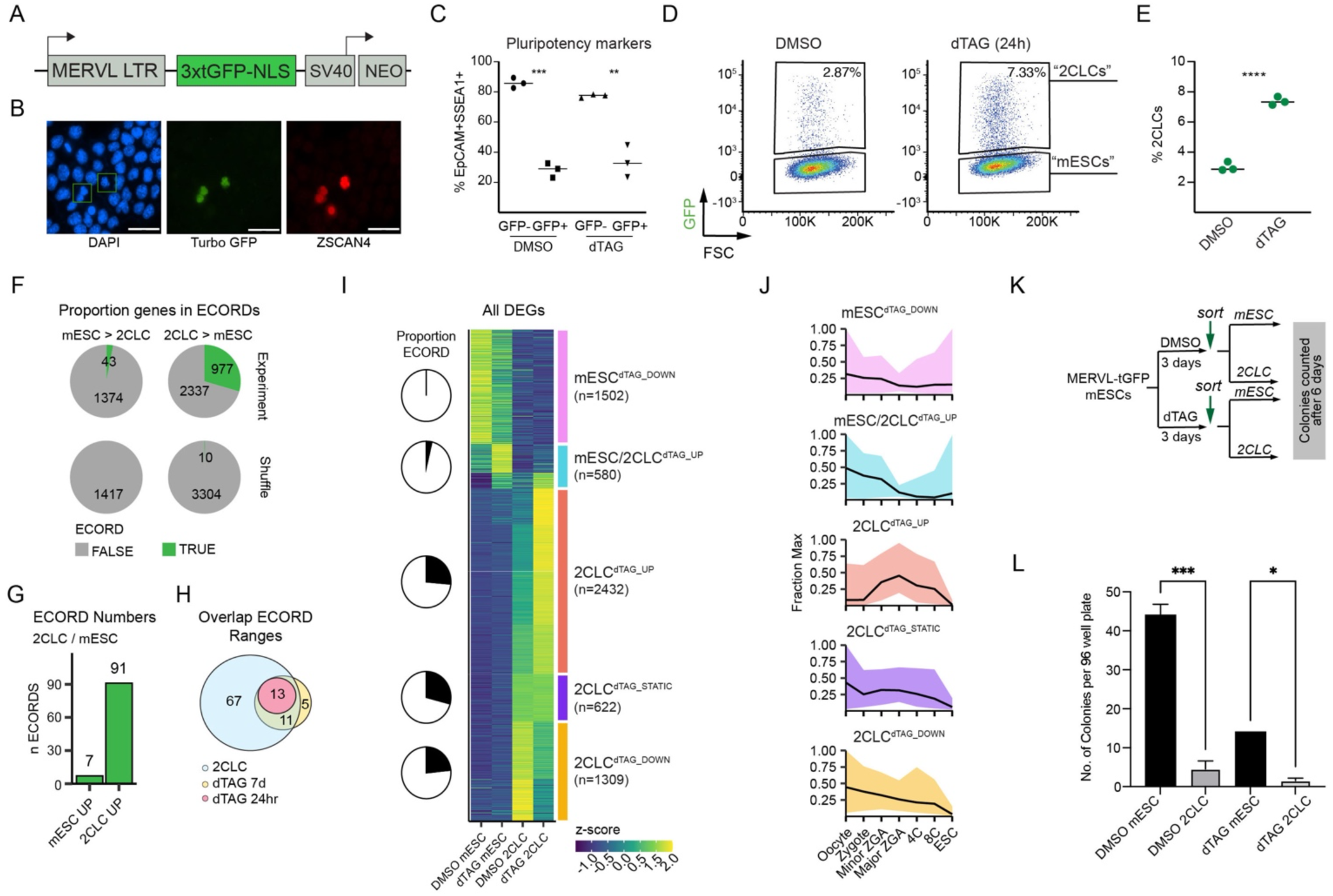
EHMT2 acts as a gatekeeper for entry into the 2-cell-like-cell state. **A)** Simplified schematic of the MERVL reporter construct used to generate MERVL-GFP mESCs. **B)** Immunofluorescence image of a representative field showing co-expression of MERVL-GFP (green) and ZSCAN4 (red). DAPI staining is indicated in blue. Green boxes indicate co-expressing cells. **C)** Percent co-expression of pluripotency-associated surface markers EpCAM and SSEA1 in mESC and 2CLC states after treatment with DMSO or dTAG, as measured by flow cytometry. **p<0.01, ***p<0.001; multiple t-tests using Holm-Sidak correction. **D**) FACS analysis of MERVL-GFP mESCs following 24h DMSO or dTAG treatment. MERVL-GFP^+^ cells are referred to as “2-cell-like-cells (2CLC)” and MERVL GFP^-^ cells as "mESCs”. **E)** Percentage of 2CLCs identified by flow cytometry after 24h dTAG or DMSO treatment (n=3 replicates). ****p<0.0001; unpaired t-test. **F)** Numbers of mESC-associated and 2CLC-associated DEGs in DMSO conditions that fall into ECORDs. **G)** Number of ECORDs in DMSO mESCs and 2CLC. **H)** Overlap of genomic ranges for ECORDs identified in bulk RNAseq (dTAG > DMSO) and ECORDs identified in sorted RNAseq (DMSO 2CLC > mESC). **I)** K-medoids clustering of all 2CLC- and mESC-associated DEGs. The signal is z-scored by row. The proportion of genes in ECORDs is indicated for each cluster. **J)** Expression (fraction max) trends of DEGs in different clusters during pre-implantation development ^30^. Only genes with an FPKM ≥10 in at least one sample type were included. The line indicates the median while the upper and lower bounds of ribbons indicate the 25 and 75 percentile. **K)** Schematic of single-cell sorting for colony-forming assays. **L)** Number of colonies formed per 96-well plate. Sorted cells were cultured in treatment media (DMSO or dTAG). *p<0.05, ***p<0.001; One way ANOVA. Error bars represent mean with SD.

EHMT2 depletion for 24h resulted in a significant increase (>2.5 fold) in the percentage of 2CLC compared to DMSO cultures (7-8% in dTAG vs 2-3% in DMSO) (**Fig.3D,E**). The increased abundance of 2CLCs in cultures treated with dTAG-13 reached statistical significance after 12h (**Fig.S3B**) and continued to grow until three days, after which the abundance stalled concomitant with cell passaging. These observations suggest that acute EHMT2 depletion in mESCs facilitates entry into the 2CLC state. Still, cells do not continue to accumulate in this state due to an apparent growth disadvantage of 2CLCs in standard mESC culture conditions.

Recent studies have revealed several distinct cellular pathways whose manipulation can increase the abundance of 2CLCs in mESC cultures, including spliceosome inhibition^33^ and retinoic acid receptor (RAR) signaling ^34^. To determine the functional interplay of these pathways with EHMT2 in modulating the mESC-to-2CLC transition, we depleted EHMT2 in MERVL-GFP mESC cultures in the presence or absence of a spliceosome inhibitor or a RAR agonist. Treatment with either compound alone significantly increased the proportion of 2CLCs (**Fig.S3D**), supporting the aforementioned prior findings. Concomitant administration of dTAG-13 further increased the percentage of 2CLCs in both instances (**Fig.S3D**), suggesting that EHMT2 activity is not affected by either treatment. We did not observe an increase in 2CLCs after treatment with two other compounds–the GSK3 inhibitor 1-Azakenpaullone and the kinase blocker WS6–used to establish cultures of cells resembling 2-cell embryos (**Fig.S3D**), suggesting that these compounds might not operate by facilitating the initial mESC-to-2CLC transition. While EHMT2 depletion and RAR activation were associated with minor changes in overall cell numbers, spliceosome inhibition significantly reduced the number of viable cells in mESC cultures (data not shown). This observation suggests that adverse selection driven by distinct metabolic requirements between mESCs and 2CLCs might partly explain the increased ratio of 2CLC cells observed upon spliceosome inhibition. In contrast, our results support that EHMT2 depletion increases the proportion of 2CLC cells in culture by facilitating the transition into the 2CLC state through a mechanism that is at least partially distinct from both retinoic acid signaling and spliceosome inhibition.

### EHMT2 has distinct gene regulatory functions in 2CLCs and mESCs

The increased abundance of 2CLCs after EHMT2 depletion could explain the apparent upregulation of ECORDs and other ZGA-associated genes we observed in RNA-seq of bulk mESC cultures. However, our analysis cannot exclude that EHMT2 depletion introduces additional transcriptional changes in 2CLCs or mESCs. In addition, bulk cell analysis likely underestimates the actual number of ECORDs in the genome. To address these limitations, we used our MERVL-GFP/EHMT2-dTAG system to perform RNA-seq on highly pure (>95%) populations of 2CLCs and mESCs three days (3d) after EHMT2 depletion (**Fig.S3E**). The comparison of DMSO-treated mESCs and 2CLCs revealed 3,314 2CLC-associated transcripts and 1,417 mESC-associated transcripts. A substantial proportion of 2CLC-associated genes (29.5%) were organized in ECORDs (977 of 3,314). In comparison, only 3% of mESC-enriched transcripts (43 of 1,417) were organized this way (**Fig.3F**). Specifically, we identified a total of 91 ECORDs in 2CLCs (mean size 10.7 genes, range 5 to 66) but only seven ECORDs in mESCs (mean size 6.1 genes; range 5 to 8) (**Fig.3G, S3F,G**)(**Table S2**). The majority of ECORDs we had detected in bulk RNA-seq (13/13 24h and 24/29 7d after EHMT2 depletion, respectively) overlapped with 2CLC-specific ECORDs (**Fig.3H)**, demonstrating that ECORDs are a feature of 2CLCs but less so of mESCs. Of note, 2CLC-associated transcripts within ECORDs showed more robust differential expression between 2CLC and mESC and higher absolute expression levels in 2CLCs compared to non-CORD genes (**Fig.S3H,I**), further supporting the strong association of ECORD activation and 2CLC identity. Of note, strong upregulation of 2CLC-associated transcripts in ECORDs, but not of transcripts outside of ECORDs, was evident in dTAG-treated mESCs (**Fig.S3J**), suggesting that de-repression of ECORDs is an early event during entry into the 2CLC state driven by EHMT2 loss. Together, these observations establish that the coordinated activation of gene clusters is a defining and widespread feature of gene expression in both spontaneously arising 2CLCs and 2CLCs triggered by EHMT2 depletion.

To further characterize the impact of EHMT2 depletion on cell state-specific gene expression, we compared the transcriptome of mESCs and 2CLCs under both DMSO and dTAG-13 conditions (**Fig.S3E**). K-medoid clustering of all genes differentially expressed in at least one pair-wise comparison (n=5,784 genes) defined five larger gene groups with distinct trends of transcriptional change in response to EHMT2 loss. Most prominently, we observed a large group of 2CLC-associated genes that were further upregulated in 2CLCs upon dTAG-13 treatment, many of them strongly ("2CLC^dTAG_UP”^) (**Fig.3I**). Smaller groups of 2CLC-associated transcripts were either weakly downregulated ("2CLC^dTAG_DOWN”^) or remained unaffected ("2CLC^dTAG_STATIC”^). All three groups of 2CLC-associated DEGs showed a similar enrichment for ECORDs (∼25% genes) (**Fig.3I**), suggesting subtle differences in the transcriptional regulation of ECORDs downstream of EHMT2 depletion. We further observed a group of mESC-associated genes that were weakly downregulated upon EHMT2 depletion ("mESC^dTAG_DOWN”^) and a group of genes upregulated in both cell types ("mESC/2CLC^dTAG_UP”^) (**Fig.3I**). Neither of these two groups showed enrichment for ECORDs, with those assigned to mESC/2CLC^dTAG_UP^ representing the rare ECORDs comprised of mESC-associated genes (**Table S2**). These results show that, in addition to facilitating the mESC-to-2CLC transition, EHMT2 depletion also significantly affects the gene expression of both mESCs and 2CLCs.

To gauge the potential biological relevance of the observation that EHMT2 depletion impacts the 2CLC gene expression program beyond facilitating the initial mESCs-to-2CLCs transition, we determined the expression kinetics of our gene groups in early mouse embryos and mESCs using published *in vivo* RNA-seq data ^30^. This revealed that 2CLC^dTAG_UP^ DEGs but neither of the other two 2CLC-associated gene groups were strongly upregulated in 2-cell embryos when ZGA occurs (**Fig.3J)**. Both groups containing DEGs associated with mESCs showed no clear association with any stage assessed, consistent with the notion that they predominantly comprise genes expressed at later stages of development (**Fig.3J**). This analysis suggests that EHMT2 depletion, in addition to facilitating entry/reprogramming of mESC into the 2CLC state, further solidifies a transcriptional program in 2CLCs that more closely resembles the developmental stage of ZGA.

2CLCs are characterized by their ability to return to a naïve pluripotent state in serum-based mESC culture conditions, though the efficiency of this process is not well-defined ^2^. To determine how the elevated expression of ZGA-associated genes in 2CLCs lacking EHMT2 might impact this process, we conducted single-cell seeding experiments with purified 2CLCs and mESCs cultured for 72h in either dTAG-13 or DMSO (**Fig.3K**). Sorted cells were allowed to grow in their respective treatment media. Quantification several days later revealed occasional mESCs colonies after seeding DMSO-treated 2CLCs, supporting the idea that these cells can revert to a pluripotent state in the presence of EHMT2. In contrast, we observed virtually no colonies with naïve pluripotent cell morphology in wells seeded with 2CLCs devoid of EHMT2 (**Fig.3L**). We also observed a reduction in the seeding efficiency of mESCs exposed to dTAG-13 compared to mESCs exposed to DMSO (**Fig.3L**), possibly reflecting impaired self-renewal of mESCs caused by increased propensity of these cells to transit into 2CLCs. This modest impairment is masked during steady mESC culture in the presence of dTAG-13. In a separate colony-forming assay, we confirmed that isolated 2CLCs that experienced EHMT2 depletion cannot revert to a pluripotent state and expand as mESCs even when cultured without dTAG-13 (**Fig.S3K,L**), likely reflecting delayed recovery of EHMT2 protein (**Fig.S2D**). Together, our observations indicate that EHMT2 loss facilitates the conversion of mESCs into 2CLCs and potentially locks these cells in that state.

### EHMT2 genome occupancy in mESCs occurs at TEs within H3K9me2 domains

Our transcriptional profiling has shown that EHMT2 represses genes both inside ECORDs (sensitive in 2CLCs) and outside of ECORDs (sensitive in 2CLCs or mESCs)(**Fig.3I**). To understand if EHMT2 regulates these genes directly, we conducted ChIP-seq experiments in EHMT2::dTAG mESCs (n=2 lines) with antibodies against the HA tag incorporated in our degron allele (see **Fig.1A**). Unlike most TFs, chromatin regulators such as EHMT2 do not directly engage with DNA, complicating reliable pulldown during ChIP-seq. Therefore, we also applied anti-HA ChIP-exo ^35^, an alternative method to map the genome occupancy of transcriptional regulators at high resolution spanning both euchromatic and heterochromatic regions ^36^. For this analysis, we implemented an alignment strategy that uses the STAR aligner to assign multi-mapping reads to the best genomic location ^37^. STAR increased the proportion of reads over repeats, particularly for ChIP-exo (**Fig. S4A**), allowing the identification of peaks that otherwise would have been missed (**Fig. S4B,C**).

In total, we detected 12,266 EHMT2 peaks with ChIP-seq and 13,675 peaks with ChIP-exo. We noticed that only a minority (∼20%) of EHMT2 peaks were called with both ChIP-seq and ChIP-exo (**Fig.S4D)(Table S4)**. To probe for evidence of EHMT2 activity, we conducted Ultra Low Input Native ChIP-seq (ULI-NCHIP) ^38,39^ against the H3K9me2 mark catalyzed by this histone methyltransferase in mESC cultures. This revealed enrichment of H3K9me2 around the EHMT2 peak summits called with either method, with the strength of signal generally correlating with the signal observed in EHMT2 ChIP (**Fig. S4D**). Together, these observations support the reliability of our data and suggest that a combination of orthogonal ChIP methods is required to detect the entire repertoire of target sites bound by chromatin regulators.

A high percentage of EHMT2 binding (∼70% for ChIP-seq and ∼90% for ChIP-exo) occurred at transposable elements (**Fig.4A**), approaching the binding frequency observed with TRIM28 (**Fig.S4E**), a key repressor of endogenous retroviruses in mESCs ^40^. EHMT2 binding to repeats significantly surpassed the numbers expected by chance or observed for pluripotency-associated transcription factors such as OCT4 and KLF4 or the reported EHMT2 recruiters WIZ ^41^ and ZFP462 ^18^ by ChIP-seq though the same pipeline (**Figs.4A, S4E**). EHMT2 binding was widespread at LINE (common and Exo-specific peaks) and LTR elements (ChIP-specific peaks) (**Fig.S4F**), suggesting a possible involvement of these elements in recruiting EHMT2 to specific gene regulatory circuits.

**Figure 4.**
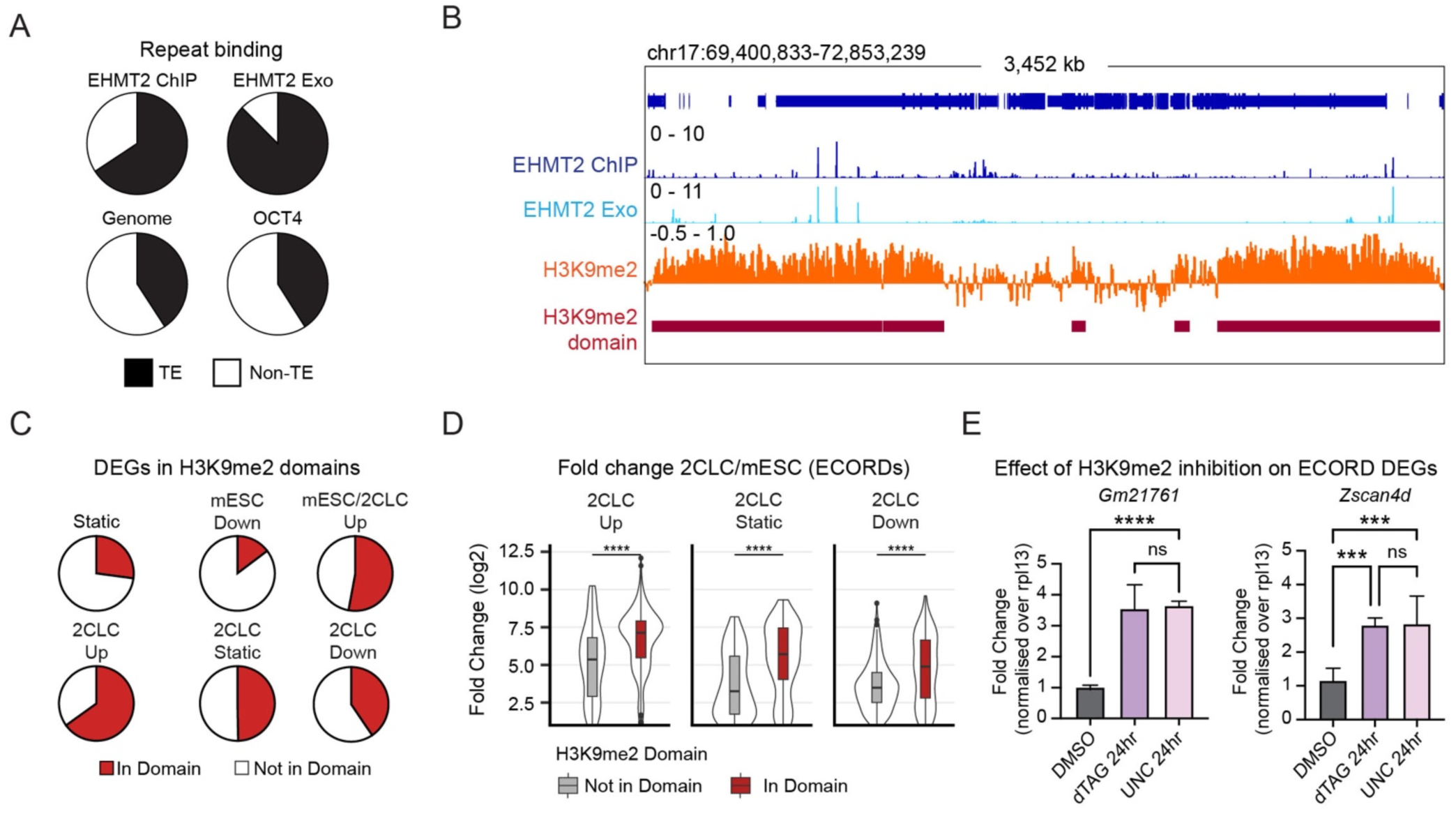
H3K9me2 domains repress ECORD expression. **A)** Proportion of indicated ChIP peaks that overlap repetitive elements. Proportion was calculated at the base pair level. **B)** Example of EHMT2 binding and H3K9me2 signal inside and outside H3K9me2 domains. **C)** Proportion of indicated DEG clusters (2CLC vs mESC) and control genes (“static”) that overlap H3K9me2 domains. **D)** Log2 Fold Changes (LFC) of DEGs in ECORDs split by whether they overlap an H3K9me2 domain. ****p< 0.0001; unpaired, two-sided, Wilcoxon Rank Sum Test with Bonferroni correction. **E)** qPCR of the ECORD genes *Gm21761* and *Zscan4d* after treatment with the H3K9me2 inhibitor UNC0638. **p<0.01, ****p<0.0001; One-way ANOVA.

EHMT2 can repress genes by nucleating heterochromatin domains that spread across the linear genome until TAD boundaries limit them ^42,43^. Consequently, individual EHMT2 binding sites do not necessarily need to overlap their transcriptional targets to exert their repressive role. To investigate the relationship between EHMT2 binding and gene expression, we therefore called H3K9me2 domains using our ULI-NChIP data, which documented that most EHMT2 binding sites were found within H3K9me2 domains (**Figs.4B, S4G**). When integrating RNA-seq and ULI-NChIP-seq data, we observed that genes upregulated upon EHMT2 depletion preferentially localized to H3K9me2 domains compared to static genes or genes that were downregulated (**Fig.4C**). This was seen both with 2CLC-associated and mESC-associated genes (**Fig.4C**) but was particularly evident for 2CLC^dTAG_UP^ DEGs in ECORDs (**Fig.S4H**). In addition, 2CLC-associated genes within H3K9me2 domains experienced more pronounced upregulation during the mESC-to-2CLC transition (**Fig.4D**) and were also upregulated upon enzymatic inhibition of EHMT2 (**Fig.4E).** These observations underscore the importance of EHMT2-regulated H3K9me2 domains for ECORD repression. Overall, these observations are consistent with the repressive nature of the observed H3K9me2 domains being dependent on EHMT2 and show that this repression is being resolved when cells enter the 2CLC state. In addition, they suggest a possible role of TEs–particularly LINEs and LTRs–in EHMT2-mediated gene repression and the establishment of H3K9me2 domains in specific genomic regions.

EHMT2 associates with different co-factors at specific categories of genomic binding sites. To gain insight into potential distinct regulatory mechanisms of EHMT2 inside and outside of ECORDs, as well as to identify possible EHMT2 co-factors or antagonists, we analyzed our combined EHMT2 ChIP peaks together with published and in-house ChIP-seq data of candidate chromatin-associated trans-acting factors (DPPA2/4, ZFP462, TRIM28, WIZ), pluripotency-associated TFs (OCT4, KLF4) and chromatin features that mark active regions of the genome (H3K27ac, ATAC-seq). We also integrated our H3K9me2 ULI-NChIP data. K-means clustering (**Fig. 5A, Table S4)** revealed four distinct categories of EHMT2 peaks predominantly defined by the mutually exclusive presence of either the zinc finger TF ZFP462, previously suggested to direct EHMT2 in mESCs towards germ layer-associated genes for repression ^18^, and the heterodimeric TFs DPPA2/4, previously shown to be required for the activation of ZGA-associated transcripts in mESCs ^3^. Thus, **k1** peaks were detected both by ChIP-seq and ChIP-exo, had strong DPPA2/4, TRIM28, and WIZ binding, strong H3K27ac and ATAC-seq signals, and strong H3K9me2 signal around the peak and surrounding genomic window. **K2** was similar to **k1** with globally lower levels of all factors and weaker H3K27ac and ATAC-seq signals but comparable H3K9me2 levels. More than 85% of k1 and k2 sites localized to H3K9me2 domains (**Fig.S5A)** and occurred almost exclusively at TEs with a striking enrichment for LINE elements (**Fig.5B**). **K3** peaks were primarily detected by ChIP-exo and showed no association with any of the tested TFs (**Fig.5A**) but were robustly marked by H3K9me2 and preferentially localized to H3K9me2 domains (**Fig.S5A**); **K4** peaks were predominantly detected by ChIP-seq, featured binding of ZFP462 and pluripotency-associated factors and exhibited strong H3K27ac and ATAC-seq signals in a pattern characteristic of active enhancers and promoters (**Fig.5A**). K4 sites were enriched for LTRs but depleted for LINEs (**Fig.5B**). Although k4 sites exhibited a local enrichment of H3K9me2 signal around the EHMT2 peak summit (**Fig.5A**), they showed only weak overlap with H3K9me2 domains relative to the genome (**Fig.S5A)**. ChromHMM analysis revealed heterochromatic features at k1/k2 and euchromatic features at k4, with k3 occupying an intermediate state (**Fig.S5B)**. Notably, weak enhancer features were seen across all peak categories (**Fig.S5B**), possibly suggesting that context-dependent activation ability is a commonality between EHMT2-bound sites. Additionally, the plurality of binding sites in all four clusters localized to gene bodies and intergenic sites (**Fig.S5C**), consistent with a predominantly promoter distal gene regulatory function of EHMT2. Overall, our k-means clustering identifies distinct categories of EHMT2 binding sites with unique chromatin features, suggesting they may have different gene regulatory functions.

**Figure 5.**
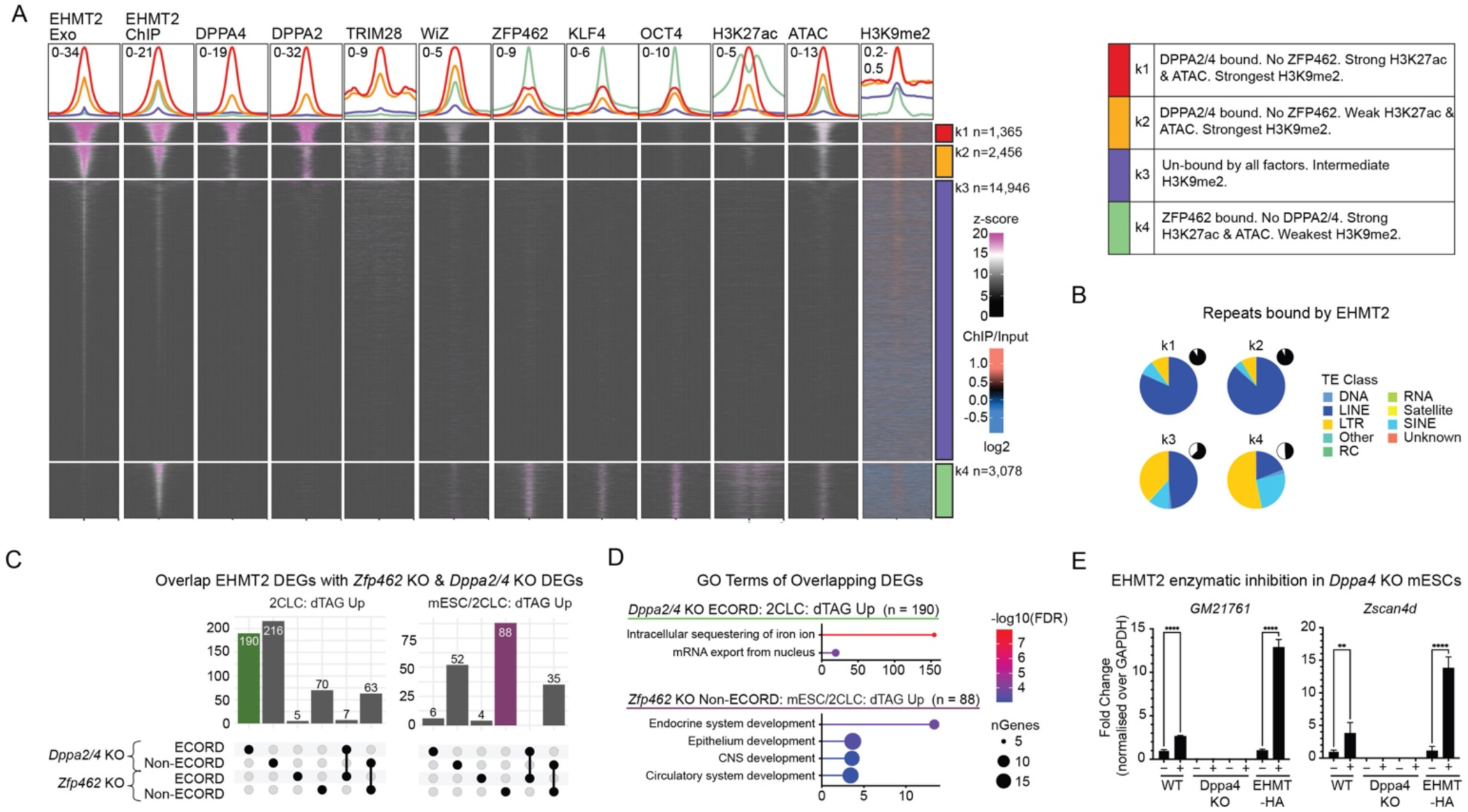
EHMT2 binding suggests locus-specific modes of gene regulation. **A)** k-means clustering of EHMT2 ChIP-Exo and ChIPseq signal in bulk mESCs with putative co-factors and factors of interest (for a list of datasets used, see Table S6). Clustering was performed on a union set of ChIP-Exo and ChIP peaks. Heatmaps are +/- 2kb from the EHMT2 peak summit, except for H3K9me2, which is +/- 50kb. For peaks called in Exo and ChIP, the midpoint of the summits was used. A summary of the properties of EHMT2 k-means clusters is included on the right. **C)** Classes of repetitive elements (TEs) that EHMT2 binds. Pie-chart insets indicate the fraction of each peak category that overlaps a repeat. **D)** Overlap of DEGs with genes <u>down</u>-regulated upon *Dppa2/4* KO (GSE126920) ^3^ or <u>up</u>-regulated upon *Zfp462* KO (GSE176321) ^18^. **E)** GO Terms of DEG categories highlighted in D. **F)** qPCR of two ECORD genes in *Dppa2/4* KO after treatment with the H3K9me2 inhibitor UNC0638. **p<0.01, ****p<0.0001; One way ANOVA. Error bars represent mean with SD.

To determine how genes transcriptionally affected by EHMT2 depletion associate with distinct EHMT2 binding modes, we integrated our ChIP categories with our RNA-seq DEGs (as defined in **Fig.3I**). This revealed a pronounced over-representation of k1/k2 (bound by DPPA2/4s) and k3 (bound by none of the tested co-factors) peaks at 2CLC^dTAG_UP^ DEGs, which was evident at gene loci both inside and outside of ECORDs (**Fig. S5D,E)**. In contrast, k4 peaks (bound by ZFP462) were overrepresented around mESC/2CLC^dTAG_UP^ genes outside ECORDs (**Fig. S5D,E)**. This suggested potentially distinct functions of DPPA2/4 and ZFP462 in regulating specific subsets of EHMT2 target loci. In line with this observation, many 2CLC^dTAG_UP^ DEGs were downregulated in DPPA2/4 KO mESCs but remained largely unaffected by KO of ZFP462 (**Fig. 5C)**. This difference was particularly pronounced for 2CLC^dTAG_UP^genes within ECORDs (**Fig. 5C)**. In contrast, mESC/2CLC^dTAG_UP^ DEGs, which predominantly localize outside of ECORDs, were more sensitive to the loss of ZFP462 and experienced upregulation in mESCs lacking this TF (**Fig. 5C)**. However, a smaller fraction of mESC/2CLC^dTAG_UP^ DEGs was downregulated in DPPA2/4 KO mESCs (**Fig. 5C),** suggesting more regulatory heterogeneity of this group of genes than for ECORDs. Functionally, 2CLC^dTAG_UP^ DEGs downregulated in DPPA2/4 KO were enriched for GO Terms related to iron and mRNA metabolism (**Fig. 5D, Table S5**). At the same time, mESC/2CLC^dTAG_UP^ DEGs upregulated upon ZFP462 KO were enriched for GO terms related to organogenesis, such as nervous and circulatory system development (**Fig. 5D, Table S5)**. These observations suggest distinct biological functions of ZFP462-controlled and DPPA2/4-controlled transcriptional programs.

The observation that the expression of ECORD DEGs depends on DPPA2/4 is consistent with EHMT2 and DPPA2/4 acting antagonistically to regulate these genes. Accordingly, enzymatic inhibition of EHMT2 in DPPA2/4 KO cells failed to result in the upregulation of ECORD DEG observed otherwise (**Fig. 5E)**, demonstrating that loss of H3K9me2 is not sufficient for de-repression of ECORDs in the absence of DPPA2/4. Together, our analyses suggest that EHMT2 regulates two broadly distinct categories of genes: 1) genes normally active later during development that in mESCs are repressed by ZFP462 acting as a co-factor for EHMT2, and 2) genes active during ZGA whose expression in 2CLCs requires DPPA2/4 and is antagonized by EHMT2. A significant fraction of the latter is organized in ECORDs.

### EHMT2 regulates the 2-cell-like state, in part, by antagonizing DPPA2/4 binding and activity

Our analysis has shown that antagonism between EHMT2-mediated repression and DPPA2/4- mediated activation is involved in determining the expression status of ECORDs and, thus, the rate of the mESC-to-2CLC transition. Since *Dppa4* is expressed in both mESCs and 2CLCs (**Fig. S6A, Table S4)**, we hypothesized that EHMT2 in mESCs might antagonize DPPA2/4 binding or activity within ECORDs. To test this model, we performed CUT&RUN ^36^ against endogenous DPPA4 in purified mESCs (MERVL-) and 2CLCs (MERVL+). We did so in dTAG-13 treated cells to obtain sufficient numbers of 2CLCs for analysis. This revealed that DPPA4 bound extensively throughout the genome (total of 42,648 sites) and was highly dynamic between the two stages, with >50% of binding sites specific to each stage (**Fig. S6B, Table S4)**. Consistent with our observations with other TFs^24^, common binding sites were enriched for promoters, while developmental stage-specific sites tended to overlap introns or intergenic sites (**Fig. S6C).** Compared to the genome-wide binding of DPPA4, which occurred primarily at sites either specific to mESCs or shared between mESCs and 2CLCs (**Fig.S6B**), 2CLC-specific DPPA4 binding sites at ECORDs were more frequent and similarly abundant to common sites **(Fig. 6A)**. This suggests that the transition from 2CLCs to mESCs—and in particular the derepression of ECORDs—may be regulated in part by the redistribution of DPPA4 to newly accessible sites.

**Figure 6.**
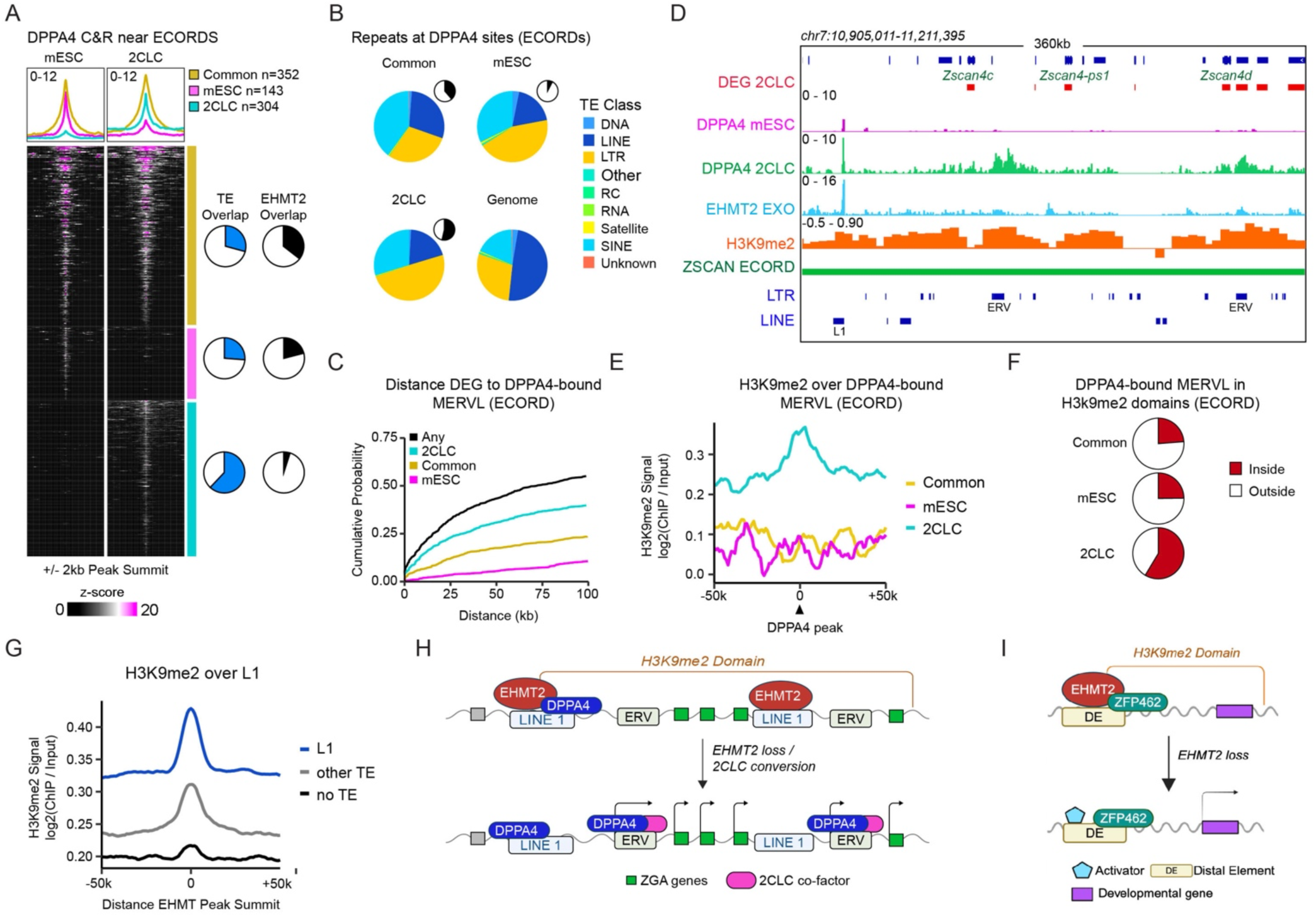
EHMT2 antagonizes DPPA2/4 function at ECORDs. **A)** DPPA4 C&R signal (+/-2kb of peak summits) around peaks associated with ECORDs. ECORD-associated peaks were identified by assigning DPPA4 peaks to the nearest expressed gene. For common peaks, the mid-point of the summits was used. **B)** Classes of repetitive elements (TEs) that DPPA4 binds. Pie chart insets indicate the fraction of all peaks that each category represents. **C)** Distance of ECORD genes to the nearest DPPA4-bound MERVL element. For this analysis, all DPPA4 peaks were used. MERVL: “ERVL” and “ERVL-MaLR” repeat families. **D)** Browser shots of EHMT2 ChIP-Exo, DPPA4 C&R, and H3K9me2 ChIP. **E)** Average H3K9me2 signal over DPPA4 MERVL within 75kb of an ECORD gene. **F)** Average signal plots of H3K9me2 signal around EHMT2 peak summits split by whether the EHMT2 peak overlaps an L1 LINE, a different type of repeat, or no repeat. **G)** Model of EHMT2 regulation of ECORDs. **H)** Model of EHMT2 regulation of Non-ECORD genes. Created with BioRender.com

What are 2CLC-specific DPPA4 binding sites? 2CLC-specific DPPA4 binding sites near ECORDs frequently overlapped TEs (**Fig.6A**) with enrichment for LTRs (**Figs.6B, S6D**) and in particular MERVL elements (**Fig.S6E**). MERVL elements are active in 2CLCs and have been suggested to act as alternative transcription start sites or enhancers ^44^ that can increase the expression of nearby genes ^2,45^. Consistent with a positive gene regulatory function in 2CLCs, 2CLC-specific DPPA4-bound MERVLs were located more proximal to ECORD DEGs when compared to common or mESC-specific sites (**Fig. 6C**). At the level of CUT&RUN tracks, we observed broad DPPA4 binding sites in 2CLCs but not mESCs near ECORD DEGs at all ECORDs analyzed **(Fig. 6D**).

We initially hypothesized that EHMT2 in mESCs might directly antagonize DPPA4 binding by occupying its binding sites and displacing DPPA4, a notion supported by the reported role of EHMT2 in suppressing MERVL expression ^20^. In contrast to these expectations, though, very few (<5%) 2CLC-specific DPPA4 sites were occupied by EHMT2 in mESCs, which argues against our hypothesis of EHMT2 directly acting on these sites (**Figs.6A, S6B**). Nevertheless, 2CLC-specific DPPA4 binding sites in mESCs were strongly marked by H3K9me2 (**Fig.6D,E**) and localized to H3K9me2 domains (**Fig.6F**). This indicates that distal EHMT2 binding may initiate the formation of H3K9me2 domains that spread through ECORDs, thereby occluding DPPA4 binding sites at MERVL proximal to 2CLC genes. We observed that EHMT2 binding sites nearest to DPPA4-bound MERVLs overlapped LINE-1 elements (**Fig.S6F**) and that EHMT2-bound LINE-1 were characterized by significantly elevated H3K9me2 levels that extended from the EHMT2 peak into their genomic vicinity (**Fig.6G**). These LINE-1 sites bound by EHMT2 localized in more gene distal positions than 2CLC-specific DPPA4 sites (**Fig.6D**) and generally comprised predominantly full-length L1Md_T/A subtypes (**Fig.S6G**). At ECORDs, EHMT2-bound L1s—but not other sites bound by EHMT2—were strongly enriched near gene-proximal MERVLs bound by DPPA4 in 2CLCs, but we notably did not find EHMT2-bound L1s enriched at 2CLC-specific DPPA4 sites nearby DEGs outside of ECORDs (**Fig. S6F**). Of note, many of these L1 were also bound by DPPA4 (**Fig.6D**). These observations suggest that EHMT2-bound L1s play a regulatory role during the silencing of ECORDs in mESCs and their reactivation during the 2CLC-to-mESC transition. In summary, we propose two distinct binding behaviors and repressive modalities of EHMT2 in mESCs. The binding of EHMT2 to gene distal LINE-1 elements in mESCs nucleates broad H3K9me2 domains that extend over clusters of 2CLC genes (ECORDs) and counteract the binding of DPPA2/4 and possibly other activating factors to proximal, LTR-derived gene regulatory elements. These H3K9me2 domains are resolved during the mESC-to-2CLC transition (**Fig.6H**). In contrast, EHMT2 represses a subset of germ layer-associated genes in mESCs through ZFP462-mediated binding to candidate enhancer elements and deposition of local H3K9me2 (**Fig.6I**).

## DISCUSSION

Unlike TFs, most chromatin regulators do not have DNA sequence specificity and, therefore, have target genes and functions that vary considerably between different cell types. We have combined acute protein depletion with different genomics assays to dissect the role of the chromatin repressor EHMT2 in naïve pluripotent stem cells. Our findings suggest that EHMT2 operates in mESCs to restrict the bi-directional differentiation capacity of these cells, counteracting both the activation of gene loci associated with “forward” germ layer differentiation and of gene loci highly expressed in 2CLCs and related to early post-fertilization development and ZGA. While the relevance of repressing the latter group of targets is underscored by the significantly increased rate of mESC-to-2CLC transitions when EHMT2 is lost, we observed no overt germ layer differentiation of EHMT2-depleted mESCs. This might reflect that EHMT2 in mESCs represses specific gene loci associated with different lineages rather than comprehensive gene expression programs. Alternatively, our observations align with redundant mechanisms of gene repression at developmental gene loci, as suggested by studies into the interaction of EHMT2 and PRC2 in mESCs ^19^.

In addition to their broadly divergent biological functions, the two groups of EHMT2 target genes we characterized differ in their genomic organization as well as the cis-regulatory elements and co-factors with which EHMT2 engages during their regulation. A significant fraction of ZGA-associated transcripts upregulated upon EHMT2 depletion are organized in what we coined ECORDs, clusters of co-regulated genes in heterochromatic regions of the genome that are poor in genes and rich in TEs. EHMT2-catalyzed H3K9me2 has been chiefly associated with gene silencing in euchromatin regions of the genome ^14,43,46^. The existence of ECORDs suggests that EHMT2 also directly contributes to developmentally controlled regions in non-euchromatic regions in mESCs, consistent with global changes in H3K9me2 levels in EHMT2 KO mESCs that are not seen in differentiated cells ^42^. ECORDs might, therefore, represent a functionally and regulatory distinct state of heterochromatin ^47^.

The activation of ECORDs in mESCs is intimately linked to the acquisition of 2CLC identity, as the observation shows that about a third of transcripts that distinguish 2CLCs from mESCs are organized in ECORDs. While activation of specific gene clusters such as ZSCAN4^28^ and OBOX ^29^ are known markers of the 2CLC stage, our findings establish clustered gene expression as a hallmark of this cell stage. Furthermore, the elevated expression of genes encoded by ECORDs during ZGA suggests that these gene clusters represent regulatory units that characterize the 2-cell state in the mouse embryo. Functionally, the organization of genes into ECORDs may enable their rapid activation and coordinated shutoff during development ^48^. In this manner, ECORDs may be loosely analogous to Hox gene clusters in that they represent clusters of genes with shared function that are coordinately deactivated through the establishment and spreading of heterochromatin domains ^49^. Supporting the importance of tight developmental control over these clusters, several ECORD-encoded genes, such as ZSCAN4^44^ and OBOX4 ^29^, have regulatory functions in 2CLCs and affect early development when inactivated.

EHMT2 controls the expression of ECORDs in antagonism with DPPA2/4, a heterodimeric pair of TFs required to express ZGA-associated genes in mESCs ^3^. Our ChIP and CUT&RUN analyses support that EHMT2 in mESCs prevents DPPA2/4 binding to MERVL-derived promoters or promoter-proximal gene regulatory elements in the context of broad H3K9me2 domains. DPPA2/4 binding to these promoters or their activation likely requires additional factors, such as DUX, which is involved in the upregulation of ZGA-associated genes during the mESC-to-2CLC transition ^5,6^.

Since we did not observe EHMT2 binding to DPPA4-bound MERVL in mESCs, it seems likely that H3K9me2 spreads from distal sites, such as local LINE-1 elements that are strongly marked by H3K9me2 and bound by EHMT2. However, we cannot rule out that our ChIP analysis missed sporadic binding of EHMT2 to MERVL. Of note, many EHMT2-bound LINE-1 are also bound by other repressors such as TRIM28 ^40^ and activators such as DPPA2/4, suggesting that these elements serve as docking sites for transcriptional regulators with diverse functions.

Control of the activation status of LINE-1 elements has been suggested to be essential for successful early mouse preimplantation development ^50^. In addition, LINE-1s have been recently proposed to serve as distal enhancers that control the expression of ZGA-associated genes such as *Zscan4* ^51^. This is consistent with the low level of H3K27ac and chromatin accessibility we have observed at EHMT2-bound k1 and k2 sites. Therefore, in addition to using LINE-1s as nucleation sites for H3K9me2 spreading, EHMT2 might counteract ECORD activation by interfering with the latent enhancer activity of these elements. Analysis of the activation status of LINE-1s in EHMT2-depleted mESCs might help to distinguish between these possibilities.

Outside of ECORDs, EHMT2 depletion resembles aspects of the transcriptional consequences of depletion of ZFP462, a sequence-specific TF, at several germ layer-associated gene loci. This agrees with a model of gene regulation in which ZFP462 recruits EHMT2 for target gene repression, as has been recently proposed for mesendodermal genes in mESCs ^18^. Our observations broadly confirm these prior observations but demonstrate that EHMT2 also represses neuroectodermal gene loci in mESCs, likely involving ZFP462. This is relevant in the context of the established role of EHMT1, the heterodimeric partner of EHMT2, in neurodevelopmental disorders ^52^. We do not rule out the existence of additional EHMT2-recruiting factors required to silence germ layer-associated genes in mESCs, such as REST^19,53^.

In summary, our experiments have provided molecular insight into the role of the chromatin repressor EHMT2 in counteracting 2CLC-specific and lineage-associated gene expression. It remains to be delineated whether the distinct modes of EHMT2 binding and target gene repression suggested by our study are also operational in other contexts—such as neurodevelopment, immune cell metabolism, and different cancers — in which EHMT2 has critical cellular functions. The identification of ECORDs, which encode many poorly characterized transcripts, also offers a novel opportunity to reveal gene regulatory aspects associated with ZGA in a tractable experimental system.

## METHODS

### Mouse cell lines

The parental mouse ESC lines used for gene targeting were KH2 ^54^ on a C57BL/6J x 129S1 F1 background. DPPA2KO, DPPA4KO, and WTJ1 cells were a kind gift from the Trono Lab ^3^.

### Mouse ESC culture

ESCs were cultured in KO DMEM (Gibco 10829018) supplemented with 15% FBS (Gemini Benchmark), 2mM Glutamax (Gibco 35050079), 0.1mM nonessential amino acids (Gibco 11140076), 100mg/ml penicillin/streptomycin (Gibco 15140163), 0.1mM 2-mercaptoethanol (Gibco 21985023) and 1000U/ml leukemia inhibitory factor prepared in house. Cells were cultured on a feeder layer of mitomycin C-treated mouse embryonic fibroblasts (MEFs) on gelatin-coated plates. In depletion experiments, cells were cultured in the presence of dTAG-13 (200nM) for indicated periods. For chemical inhibition of EHMT2, cells were cultured in mESC media with UNC0638 (1μM). Cells cultured in DMSO served as controls.

### Generation of EHMT2-dTAG mESCs

To generate EHMT2-dTAG mESCs, homology arms covering about 1.2kb of sequence around the *Ehmt2* stop codon were PCR-amplified from KH2 genomic DNA and cloned into pBluescript (Stratagene) vector together with an FKBP12F36V-2xHA-P2A-NLS-mCherry cassette, using Gibson assembly. Parental KH2 mESCs were co-transfected with the targeting vector and a pX330-neo^R^ vector expressing Cas9 and gRNAs targeting *Ehmt2* using TransIT-293 (Mirus Bio 2700). The next day, cells were plated on a 10cm plate at low density and cultured in selection media containing Geneticin (500mg/ml) for 48 hours. Individual clones were picked, expanded, and confirmed using PCR, Sanger sequencing of PCR amplicons, and flow cytometry. Guide RNAs are listed in Supplemental Table S6.

### Generation of MERVL-EGFP mESCs

EHMT2-dTAG mESCs were transfected with the 2C-3XtbGFP-PEST plasmid ^31^ (Addgene #69072) using lipofection with TransIT-293 (Mirus Bio 2700). Cells were seeded at clonal density and selection in G418 (250 mg/ml) for five days before the cultures were inspected under an EVOS fluorescence microscope, and individual colonies with rare GFP^+^ cells were picked for further expansion.

### Western blotting

Three independent clones were treated with DMSO or dTAG for 24 hours before protein isolation. Cells were washed with PBS -/- and harvested using Trypsin (Life Technologies 25200114). Nuclear lysates were prepared using the NE-PER kit (Thermo Fisher 78835) according to the manufacturer’s instructions. Histones were isolated using the histone extraction kit (Abcam ab113476) according to the manufacturer’s instructions. Protein concentration was measured using Bradford Reagent (BioRad 5000201), and samples were boiled in Laemmli Sample Buffer (BioRad) with beta-mercaptoethanol and run on Invitrogen precast gels. Blots were imaged using Azure Biosystems C400. Images were quantified using ImageJ ^55^. The following antibodies were used at 1:1000: anti-HA (Abcam 9110), anti-H3K9me2 (ab1220), and anti-histone H3 (Abcam 1791).

### Flow cytometry

Expression of mCherry in EHMT2-dTAG mESCs was determined using the 561nm (610/20) channel on a BD Fortessa. For the EHMT2 and H3K9me2 depletion kinetics cells, two independent clones were treated with DMSO or dTAG (200nM) for the indicated periods. 500k cells from each treatment group (technical triplicates) were collected in a single-cell suspension using a Thermo Fisher intracellular staining kit (cat. 00-5523-00). Cells were fixed for 25 min and incubated with 50ul primary antibodies diluted in permeabilization buffer for HA (CST-3724S 1:800) and H3K9me2 (ab1220 1:400) for 30min. Cells were then incubated with 50ul secondary antibodies at 1:500 dilution (A-31572 555aRb, A-31571 647aMs) for 30 minutes in the dark. Samples were run through the BD Fortessa flow cytometer and analyzed using the FloJo software. Significance was called using the R t_test default function in the rstatix package ^56^. The percentages of mESCs expressing the MERVL-EGFP reporter and the pluripotency-associated surface markers SSEA-1 and EpCAM were determined using live-cell flow cytometry using a BD FACSCanto. Similarly, cell-sorting of the MERVL-GFP+ and MERVL-GFP- was conducted with the help of the flow cytometry core facility at Weill Cornell. Live cells were run on the BD Influx on the FITC channel and gated based on high and low fluorescence, followed by cell sorting. Data analysis was done using FloJo software.

### UNC0638 treatment and RT-qPCR

EHMT2-dTAG mESCs were treated with DMSO, dTAG, or UNC0638 inhibitor for 24 hours. RNA was isolated from treated samples, as mentioned above. Reverse transcription of RNA from each sample was performed using the iScript kit (BioRad 1708841). qPCR was performed on cDNA samples in triplicate using PowerUp SYBR green PCR master mix (Thermo Fisher A25778) on an Applied Biosystems QuantStudio3. The primers used are in Table S6.

### RNA isolation

Total RNA from cells was extracted using TRIzol (Invitrogen 15596018) and purified using the RNA Clean and Concentrator kit (Zymo Research ZR1014). RNA quality and quantity were checked before assays using a nanodrop or bioanalyzer.

### RNA Sequencing

For the recovery experiment, EHMT-dTAG mESC clones (triplicates) were treated with DMSO or dTAG (200nM) for 24 hours, 7 days, and 15 days. After 7 days of dTAG treatment, the recovery group was allowed to grow in DMSO for 8 days. Cells were collected at the end of the treatment cycle, and RNA was extracted as described above. Using the manufacturer’s instructions, 1ug of RNA was then used to make libraries using the TruSeq Stranded mRNA Library Prep (Illumina# 20020595). Libraries were sequenced on the NovaSeq 6000 using the Weill Cornell genomics core’s S4 flow cell at PE 2X100. For the MERVL reporter experiments, MERVL-GFP mESCs were treated in triplicate with DMSO or dTAG (200nM) for 72 hours and sorted into GFP^-^ and GFP^+^ populations, as described in flow cytometry. 200,000 cells were collected, and RNA was isolated using TriZol, as described above. Low-input RNA libraries were prepped by Novogene and sequenced libraries at PE x150 on a Novaseq 6000.

### Immunofluorescence

EHMT-dTAG-MERVL cells treated with DMSO or dTAG for 24hr were washed and fixed in 4% formaldehyde for 10mins, blocked in blocking buffer (PBS-/- with 1% BSA, 0.1% Triton-X-100, and 3% donkey serum (Sigma D9663) and stained with ZSCAN4 antibody (AB4340) at 1:150 for 2hrs. Cells were washed thrice with PBS -/- with 0.1% Triton X-100 (PBST) and then incubated in Donkey anti-Rabbit Alexa Fluor 555 secondary antibody at 1:1000 for 1 hour. Cells were washed thrice with PBST. DAPI (300nM) was added for 5 minutes for nuclear staining. Images were taken on a Nikon fluorescent microscope. Images were quantified using ImageJ.

### Colony Forming Assay

EHMT-dTAG-MERVL cells were treated with DMSO or dTAG for 72h and sorted by flow cytometry as described above into GFP+ (2CLC) and GFP- (mESC) populations. Single cells were seeded into two 96-well plates per treatment group. Cells were allowed to grow in mESC media for 6 days. The number of colonies per plate was then counted. Alternatively, 1000 sorted cells (2CLC and mESC) were plated into each well of a 6-well plate and allowed to grow in either DMSO or dTAG. Cells were allowed to grow for 5 days, after which alkaline phosphatase staining was done using Vector® Red Alkaline Phosphatase Substrate Kit (SK-5100) using the manufacturer’s instructions, and images were taken. The number of colonies was counted using ImageJ.

### ATAC-Seq

ATAC-seq was performed as previously described ^57^ with some modifications. Briefly, cells from two independent clones (EHMT2-dTAG mESCs) were treated with DMSO for 24hrs. Cells were trypsinized, and 50,000 cells per replicate were washed with 50 μL cold 1xPBS followed by 50 μL lysis buffer (10 mM Tris-HCl pH 7.4, 3 mM MgCl2, 10 mM NaCl, 0.2% (v/v) IGEPAL CA-630) to isolate nuclei. Nuclei were pelleted by centrifuging for 10min at 800g at 4°C and 50μL transposition reaction mix (25 μL TD buffer, 2.5 μL Tn5 transposase, and 22.5 μL ddH2O) was added. Reagents from the Nextera DNA library Preparation Kit (Illumina #FC-121–103) were used. Samples were incubated at 37°C for 30min. DNA was isolated using the ZYMO Kit (D4014). ATAC-seq libraries were generated using NEBNext High-Fidelity 2X PCR Master Mix (NEB, #M0541), with each sample assigned a unique barcode and a universal primer. The optimal cycle number for each sample was determined by qPCR. Samples were size-selected (0.55x–1.5x) using SPRIselect beads (Beckman Coulter, B23317). Libraries were assessed with an Agilent Bioanalyzer. Libraries were sequenced on an Illumina Nova-Seq 6000 platform with 100bp paired-end reads.

### ChIP-Seq

Two independent clones (EHMT2-dTAG mESCs) were treated with DMSO 24 hours before ChIP-seq. ChIP was performed as previously described ^58^ with some modifications. Briefly, 30 million cells per replicate for each condition were double-crosslinked by first incubating with 2mM DSG (Sigma 80424-5MG-F) for 50 mins, followed by 1% formaldehyde at RT for 10 minutes, and then quenched with 125mM glycine for 5 mins at RT. Cells were resuspended in 300ml lysis buffer (10mM Tris pH8, 1mM EDTA, 0.5% SDS) and sonicated in a Pico bioruptor for 15-30 cycles and then centrifuged for 10 minutes at 4°C at 17000g. Supernatants were precleared for 1hr with 20ul of protein A Dynabeads (ThermoScientific 10-001-D) per sample. 5% of each sample was removed and frozen as an input sample. Samples were diluted 5 times with dilution buffer (0.01% SDS, 1.1% triton,1.2mM EDTA,16.7mM Tris pH8, 167mM NaCl) and incubated with HA antibody (5ug/30M total cells) (CST 3724) O/N with rotation at 4°C. The next day, protein A Dynabeads pre-blocked with 1mg/ml BSA protein were added to each sample (30ul Dynabeads per sample). Samples were incubated for 3.5 hours at 4°C. Subsequent steps were carried out as previously described. According to the manufacturer’s instructions, 15ng of immunoprecipitated DNA and input were used for library amplification with the KAPA Hyper prep kit (KK8502). Libraries were sequenced on an Illumina Novaseq 6000 platform with 100bp paired-end reads.

### ChIP-EXO

Cultures of EHMT2-dTAG mESCs were treated with DMSO for 24 hours. Cells were collected by trypsinization for 3 mins and were washed with PBS, followed by centrifugation (500 g, 5 mins, 4°C). 10M cells were used per replicate. The pellet was resuspended to have 1 million cells per mL in PBS (RT). Cells were crosslinked with 1% formaldehyde in RT for 10 minutes on a platform shaker. Crosslinking was quenched with 125 mM glycine for 5 min at RT on a platform shaker. Crosslinked cells were centrifuged (500 g, 5 mins, 4°C) and were washed twice with ice-cold PBS. After the final wash and centrifuge, the pellet was snap-frozen before extraction. Frozen cell pellets were processed as described previously in the ChIP-exo 5.0 protocol ^35^. Anti-HA antibody (Abcam 9110) was used for o/n chromatin immunoprecipitation at 4°C. Immunoprecipitated material was processed as described ^35^.

### ULI-NChIP

ULI-NChIP was conducted following a published procedure ^38^, using 100,000 cells as input for the bulk ULI-NChIP with two biological replicates. Briefly, frozen cell pellets were resuspended in Sigma EZ nuclei isolation lysis buffer (NUC101) with 1x protease inhibitor cocktail (Roche 04693132001) and 1mM PMSF (TF 36978) and digested with MNase (NEB M0247) as suggested by the protocol. The digested chromatin was rotated for 1 hour at 4C, followed by pre-clearing using Protein A/G beads (Dynabeads, Life Technologies #1006D), and 10% then saved as input control. 1ug per 100k cells of H3K9me2 antibody (ab1220) was added to Protein A/G beads for 3h at 4C to form the antibody complex. Precleared chromatin was added to the antibody complex and rotated at 4C overnight. The antibody-bound chromatin was then eluted from the magnetic beads using 100mM NaHCO3 and 1%SDS at 65C for 1h and purified alongside the input chromatin. Libraries of the eluted and input chromatin were prepared using the Kapa Hyperprep kit (KK8502) following the manufacturer’s instructions. Libraries were sequenced on an Illumina NovaSeqXplus platform with 100bp paired-end reads.

### CUT&RUN

MERVL-GFP mESCs (in duplicates) were treated with dTAG-13 for 72 hours and sorted based on GFP as mentioned above in flow cytometry. CUT&RUN against DPPA4 was performed on 17000k-50000 sorted cells, as previously described^59,60^. Live cells were sorted based on GFP fluorescence on an Influx Cell Sorter before performing CUT&RUN. BioMag Plus Concanavalin A beads (ConA beads, PolySciences NC1358578) were washed twice with cold bead activation buffer (20mM HEPES pH7.5, 1mM MnCl_2,_ 10mM KCl, 1mM CaCl_2_,). Sorted cells were washed twice in wash buffer (20mM HEPES pH7.5, 150mM NaCl, 0.5mM Spermidine (Acros AC132740050), ½ Protease Inhibitor tablet). At room temperature, 20 ml of activated ConA beads per sample were bound to the cells in wash buffer for 10 minutes. Samples were incubated overnight with gentle rocking at 4°C with 1:100 of anti-DPPA4 (AF3730) or 1:100 anti-IgG (EpiCypher 130042) in antibody buffer (Wash Buffer with 2mM EDTA and 0.01% Digitonin (Millipore Sigma 300410)). Samples were washed twice in Digitonin Buffer (Wash Buffer with 0.01% Digitonin). ProteinA/G MNase (EpiCypher) was then bound to samples for 1 hour at 4°C. ProteinA/G MNase was diluted in 50ul antibody buffer. Samples were washed twice in Digitonin Buffer, and 1ml 100mM CaCl_2_ was added to activate MNase digestion. After 2hrs of incubation, MNase digestion was quenched using 33ml STOP Buffer (340mM NaCl, 20mM EDTA, 4mM EGTA, 50mg/ml RNAse A, 50mg/ml glycogen, 0.015 ng/ml E. coli spike-in) and samples incubated at 37°C for 20 minutes. Digested fragments were isolated by centrifuging samples for 5 minutes at 16,000g, binding ConA beads to a magnet for 2m, and then saving the supernatant. DNA was purified by adding 0.1% SDS and 5ug Proteinase K for 10 minutes at 70°C, followed by phenol-chloroform extraction and precipitation in 100% ethanol at −80°C overnight. Pellets were washed in 100% ethanol and resuspended in 12ml nuclease-free water. CUT&RUN libraries were prepared using the ThruPLEX DNA-Seq (Takara, Cat #: R400675) using and Unique Dual Index (Takara #R400665) kits according to the manufacturer’s instructions until the amplification step. After adding indexes, amplification cycles were performed using a shortened annealing/extension time (67C, 10s) to enrich for small fragments, as previously recommended ^59^. Libraries were size selected with 1.5x volume SPRI beads (Beckman Coulter B23317) and sequenced at the Weill Cornell Genomics Resources Core Facility on an Illumina NovaSeq 6000 (PE-100, 30 million reads per sample).

### Data Processing and Analysis

All genomics datasets were processed using custom snakemake (v6.6.1) pipelines (31).

### RNA-seq

#### Initial processing and explanation of STAR multi command

Technical replicates were merged using zcat. Reads were trimmed to remove adapter sequences using trim galore (v0.6.7)^61,62^ with parameters: --phred33 --quality 0 --stringency 10 --length 20. Trimmed reads were mapped using STAR (v2.7.10)^63^. STAR alignment was performed using a previously described strategy developed to map multi-mapping reads to the best location in the genome (hereafter, “STAR multi”)^37^ using parameters: --readFilesCommand zcat --runThreadN 10 --outSAMtype BAM SortedByCoordinate --outFilterMultimapNmax 5000 - -outSAMmultNmax 1 --outFilterMismatchNmax 3 --outMultimapperOrder Random -- winAnchorMultimapNmax 5000 --alignEndsType EndToEnd --alignIntronMax 1 -- alignMatesGapMax 350 --seedSearchStartLmax 30 --alignTranscriptsPerReadNmax 30000 -- alignWindowsPerReadNmax 30000 --alignTranscriptsPerWindowNmax 300 -- seedPerReadNmax 3000 --seedPerWindowNmax 300 --seedNoneLociPerWindow 1000 -- alignSJoverhangMin 999 --alignSJDBoverhangMin 999. A detailed description of the modified STAR command is provided below. TE Counts (v2.2.3)^64^ was used to assign mapped reads to protein-coding and repeat genes and repeats with default parameters. GTFs from GENCODE (vM25)^65^ and a repeat file from TE Counts were used. Bigwigs were made from bed files using deeptools (v3.5.1)^66^ using parameters: -p 10 --binSize 10 --ignoreForNormalization chrX chrM - -normalizeUsing RPGC --effectiveGenomeSize 2652783500 --extendReads 200 – ignoreDuplicates.

| Explanation of STAR multi Command |  |  |
| --- | --- | --- |
| Flag | Value | Description |
| outFilterMultimapNmax | 5000 | The max number of places in the genome STAR will allow a read to align. Default is 10. |
| outSAMmultNmax | 1 | The number of entries STAR will write for each multi-mapper. Top scoring alignments are outputted first. |
| outFilterMismatchNmax | 3 | Number of mismatches to allow to be outputted. Default is 10 |
| outMultimapperOrder | Random | STAR should output the order of multimapping reads in random order. Without runRNGseed set alignments will vary between runs. |
| winAnchorMultimapNmax | 5000 | The number of loci that anchors can map to. winAnchorMultimapNmax must be >= outFilterMultimapNmax. Default is 50. |
| alignEndsType | EndToEnd | Turn off soft-clipping of reads. |
| alignIntronMax | 1 | Maximum intron size. Default is 0. |
| alignMatesGapMax | 350 | Maximum gap between two mates. Default is 0. |
| seedSearchStartLmax | 30 | Search start point through the read. The read is split into pieces <= this value. Default is 50. |
| alignTranscriptsPerReadNmax | 30000 | Max number of different alignments per read to consider. Default is 100 |
| alignWindowsPerReadNmax | 30000 | The max number of windows per read. Default is 100 |
| alignTranscriptsPerWindowNmax | 300 | The maximum number of transcripts per window. Default is 100 |
| seedPerReadNmax | 3000 | Max number of seeds per read. Default is 1000. |
| seedPerWindowNmax | 300 | Max number of seeds per window. Default is 50. |
| seedNoneLociPerWindow | 1000 | Max number of one seed loci per window. Default is 10 |
| alignSJoverhangMin | 999 | Minimum overhang for spliced alignments. Default is 5 |
| alignSJDBoverhangMin | 999 | Minimum overhang for annotated junctions. Default is 1 |

#### Analysis

DESeq2 was used to identify differentially expressed genes ^67^. Differentially expressed genes were defined as genes with an adjusted p-value <0.05 and log2 fold-change (LFC) ≥1.

Expressed genes were defined as genes that remained after DESeq2 performed independent filtering to remove genes with low counts (pAdj == “NA”). Normalized counts were generated using the counts function in DESeq2. For k-medoid clustering, normalized counts were z-scored, and a distance matrix was generated using the distNumeric function from the kmed package^68^ with Manhattan weighted range (mrw). All genes were first assigned to a TAD to identify clusters of genes for ECORD analysis ^24^. Gene lists were then filtered to all differentially expressed genes, and the number of DEGs that occurred in sequence along the linear genome was counted. Clusters were broken if: 1) the next DEG was changed in the opposite direction (p-adj < 0.05, no fold-cutoff), 2) the next gene was in a different TAD. After assigning each gene a cluster ID, the number of static genes was calculated by taking the full range of each cluster and intersecting it with the ranges of static genes. For subsequent analyses, “ECORDs” were defined as clusters with ≤50% static genes as well as ≥4 DEGS (LFC > 1) for the bulk dataset, and ≤50% static genes and ≥5 DEGs (LFC>1) for the sorted-cell RNAseq dataset. For ChromHMM, gene promoters (TSS −2.5kb/+0.5kb) were used with a published, 100-state model ^26^. After running, the values were re-scaled from 0-1 per column. For compartment analysis, the mean compartment signal across the entire gene body was calculated using a published dataset ^24^. For comparison to *in vivo* embryonic development, we used published FPKM ^30^ and gene categories from Hu et al.^69^. To correct gene.ids in the *in vivo* dataset that did not match GENCODE annotations, the ‘select’ function was used to obtain aliases from the org.Mm.eg.db package, which was sufficient to correct nearly all inconsistencies. All plots were generated with ggplot2 ^70^. For comparison to Zfp462 and Dppa4 DEGs, published datasets were reprocessed, and DEGs were called using the methods described above. The following published datasets were used: GSE176321 (zfp462 cl1, zfp426 cl2, WT ESC), GSE126920 (Dppa2 KO, Dppa4 KO, WT ESC).

### Gene Ontology (GO) analysis

Gene ontology analysis was done using ShinyGO ^71^. Lists of genes were uploaded on the ShinyGO v0.81 server with the following settings: ‘Mouse’ as the ‘Best matching species’, ‘0.05’ as the ‘*P* value cutoff (FDR)’, and ‘25’ as the ‘# of most significant terms to show’. All expressed genes (see above) in each dataset were used as background. For 24h ECORD and non-ECORD, the terms obtained were collapsed using rrvgo (v3.19)^72^ and represented using R. For GO terms of overlapping DEGs, the most relevant terms were selected and represented using R.

### ChIPseq and ChIP-Exo

#### Initial Processing

Reads were concatenated, trimmed, and aligned as for RNAseq above. PCR duplicates were removed using Picard (v2.26.0)^73^ with parameters: VALIDATION_STRINGENCY=LENIENT REMOVE_DUPLICATES=true ASSUME_SORTED=true. After duplicate removal, reads that overlapped ranges annotated as "High Signal Region" from the ENCODE Blacklist (mm10, v2) were removed using samtools (v1.14) ^74^. Bam files were converted to bedpe files using bedtools (v2.30.0) ^75^ with parameters: -bedpe and then reformatted using the command: “sort -k 1,1 -k 2,2n | cut -f 1,2,6,7”. Peaks were called on bed files using MACS2 (v2.2.6) ^76^ using parameters: -f BEDPE --nomodel --seed 123 --keep-dup all -p 0.05. After peak calling with MACS2, peak files were filtered only to include peaks with q-value < 0.05 using a custom Python script. Bigwigs were made from bed files using deeptools (v3.5.1) ^66^ using parameters: --binSize 10 -- ignoreForNormalization chrX chrM --normalizeUsing RPGC --effectiveGenomeSize 2652783500 --extendReads 200 --ignoreDuplicates --blackListFileName mm10-blacklist.v2_highSignalRegions.bed. Bigwigs were z-normalized per chromosome arm using a custom R script (v4.1.2). The following published datasets were also used: GSE137272 (Wiz), GSE158460 (Trim28), GSE177058 (Zfp462), GSE74112 (Oct4), GSE95517 (Dux), GSE113429 (H3K27ac, Klf4), GSE126921 (Dppa2, Dppa4).

ULI-NChIP reads were processed identically to ChIPseq reads through to the generation of bigwigs. After generating RPGC normalized bigwigs, the bigwigs were re-binned at 10kb resolution with a 5kb sliding step using a custom R script as previously described ^77^. Briefly, the mm10 genome was tiled into non-overlapping 5kb bins. The bins were re-sized to 10kb, anchoring at the center, and the mean RPGC normalized signal across the 10kb window was calculated. Locations with 0 signal were ignored as these likely represent regions of the genome annotated as “High Signal Regions,” which were removed during processing or unmappable regions. Bins were then resized to 5kb before export as a bigwig. Consequently, each bin in the bigwig represents the mean across the 5kb window +/- 2.5kb. Input normalized tracks were generated by calculating log2(ChIP/Input) per bin. To identify H3K9me2 domains, regions of the genome with high H3K9me2 signal (log2(ChIP/Input) > 0.5, ∼75 percentile) were extended up- and down-stream until a region without H3K9me2 enrichment was encountered (log2(ChIP/Input) < 0). The following packages were used: Genomic Ranges ^78^, plyranges ^79^, rtracklayer ^80^, and org.Mm.eg.db ^81^.

#### Analysis

For repeat analysis, the mm10_rmsk_TE.gtf repeat masker file from TE Transcripts ^82^ was used. The fraction of binding sites that overlap repeats was done at the base-pair level by calculating the number of bases within each peak that overlapped a repeat. This was done to make the values comparable to the whole genome fraction. For k-means clustering, union ranges of EHMT ChIP-seq and ChIP-Exo were generated, and the locations of the peak summits were annotated. The summit from the respective dataset was used for peaks called in specific to either ChIP-seq or ChIP-exo. For common peaks, the midpoint between the two summits was used. The signal from z-normalized bigwigs +/- 250bp of peaks summits for all datasets, except for H3K9me2, was obtained using the getPlotSetArray function from seqplots ^83^, updated to make it compatible with Bioconductor >v3.14 (https://github.com/cmuyehara/seqplots), using parameters: refgenome = ‘mm10’, bin = 10, rm0 = F, ignore_strand = T, xanchored = ‘mf’, add_heatmap = T. For H3K9me2 ULI-NChIP, +/-50k signal from the input-normalized bigwigs was used for clustering with the seqplots parameter modified to use a 1000bp bin. Heatmaps from all datasets were clustered together using the kmeans function in R after running set.seed(883). ChromHMM analysis on k-means clusters was performed on union peak ranges as for RNAseq above. To calculate the genomic distribution of binding sites in k-means clusters, the annotatePeakInBatch function from ChIPpeakAnno was used with parameters: PeakLocForDistance = ‘middle’, output = ‘shortestDistance’, FeatureLocForDistance = ‘middle’, ignore.strand = T. To associate EHMT peaks with RNAseq, each expressed gene was assigned to the nearest EHMT peak (from the union dataset) in the same H3K9me2 domain. To calculate enrichment, a bootstrap approach was used in which gene categories (2CLC > mESC, static, etc.) were shuffled 1,000 times.

### CUT&RUN

#### Initial Processing

Technical replicates were merged using zcat. Reads were trimmed using the bbduk.sh command from BBMap ^84^ using parameters: ktrim=r ref=adapters rcomp=t tpe=t tbo=t hdist=1 mink=11. Trimmed reads were aligned using STAR as for RNAseq above. Duplicate and blacklist removal were performed as for ChIPseq above. Peaks were called on bed files using MACS2 (v2.2.6) ^76^ using parameters: -f BEDPE --nomodel --seed 123 --keep-dup all. Bigwigs were z-normalized per chromosome arm using a custom R script.

#### Analysis

Analysis was performed as described above for ChIPseq and ChIP-Exo.

### ATACseq

ATAC-seq was processed similarly to CUT&RUN above. To correct for the tn5 cut position, fragments were shifted +4bp and −5bp for reads mapping to the + and - strand, respectively, using a custom awk script as recommended ^85^.

### Quantification and statistical analyses

Statistical analysis of western blot, flow cytometry, and colony assays was done on PRISM (GraphPad Prism version 10.0.0, GraphPad Software, Boston, Massachusetts, USA, www.graphpad.com). Specific tests and corrections applied are indicated in the respective figure legends and supplemental table S7.

Table of reagents and antibodies.

| Antibodies | Source | Identifier |
| --- | --- | --- |
| HA | Cell signalling technology, Abcam | CST 3724S, ab9110 |
| H3K9me2 | Abcam | ab1220 |
| SSEA1, Biotin-conjugated | Thermo Fisher | 13-8813-82 |
| EpCAM (CD326), PECy7-conjugated | Thermo Fisher | 25-5791-80 |
| ZSCAN | Millipore-Sigma | AB4340 |
| DPPA4 | Novus | AF3730 |
| Donkey anti-Rabbit Alexa Fluor 555 | Thermo Fisher Scientific | A-31572 |
| Donkey anti-Mouse Alexa Fluor 647 | Thermo Fisher Scientific | A-31571 |
| **Chemicals** |  |  |
| DSG (Di(N-succinimidyl) glutarate) | Sigma | 80424-5MG-F |
| dTAG |  |  |
| UNC0638 | Selleckchem | S8071 |
| **Commercial assays** |  |  |
| eBioscience™ Foxp3 / Transcription Factor Staining Buffer Set | Thermo Fisher Scientific | 00-5523-00 |
| KAPA hyper prep-kit | Roche | KK8502 |
| **Software and algorithms** |  |  |
| snakemake | 86 | https://anaconda.org/bioconda/snakemake |
| Trim Galore | 61,62 | https://anaconda.org/bioconda/trim-galore |
| STAR | 87 | https://anaconda.org/bioconda/star |
| samtools | 74 | https://anaconda.org/bioconda/samtools |
| Picard | 73 | https://anaconda.org/bioconda/picard |
| bedtools | 75 | https://anaconda.org/bioconda/bedtools |
| BBTools | 84 | https://anaconda.org/bioconda/bbmap |
| Deeptools | 66 | https://anaconda.org/bioconda/deeptools |
| TE Transcripts | 64 | https://anaconda.org/bioconda/tetranscripts |
| MACS2 | 76 | https://anaconda.org/bioconda/macs2 |
| org.Mm.eg.db | 81 | https://bioconductor.org/packages/release/data/annotation/html/org.Mm.eg.db.html |
| rtracklayer | 80 | https://bioconductor.org/packages/release/bioc/html/rtracklayer.html |
| GenomicRanges | 78 | https://bioconductor.org/packages/release/bioc/html/GenomicRanges.html |
| GenomicFeatures | 78 | https://bioconductor.org/packages/release/bioc/html/GenomicFeatures.html |
| plyranges | 79 | https://bioconductor.org/packages/release/bioc/html/plyranges.html |
| seqplots | 83 | https://github.com/cmuyehara/seqplots |
| DESeq2 | 67 | https://bioconductor.org/packages/release/bioc/html/DESeq2.html |
| ChIPpeakAnno | 88 | https://bioconductor.org/packages/release/bioc/html/ChIPpeakAnno.html |
| kmed | 68 | https://cran.r-project.org/web/packages/kmed/index.html |
| tidyverse | 89 | https://cran.r-project.org/web/packages/tidyverse/index.html |
| ggplot2 | 70 | https://cran.r-project.org/web/packages/ggplot2/index.html |
| patchwork | 90 | https://cran.r-project.org/web/packages/patchwork/index.html |
| eulerr | 91 | https://cran.r-project.org/web/packages/eulerr/index.html |
| ComplexUpset | 92 | https://cran.r-project.org/web/packages/ComplexUpset/index.html |
| fishualize | 93 | https://cran.r-project.org/web/packages/fishualize/index.html |
| ggalluvial | 94 | https://cran.r-project.org/web/packages/ggalluvial/index.html |
| rstatix | 56 | https://cran.r-project.org/web/packages/rstatix/index.html |
| ImageJ | 55 | https://imagej.net/ij/ |
| rrvgo | 72 | https://www.bioconductor.org/packages/release/bioc/html/rrvgo.html |

## RESOURCE AVAILABILITY

### Lead contact

Requests for further information, resources, and reagents should be directed to and will be fulfilled by the lead contact, Matthias Stadtfeld.

### Materials available

Cell lines generated in this study are available upon request from the lead contact.

### Data and code availability

RNA-seq, CUT&RUN, ChIP-seq, ChIP-exo, and ULI-NChIP-seq data have been deposited at Gene Expression Omnibus (GEO) under accession code GSE280606 (<u>Reviewer Token</u>: yvyxoauunfkzdcn). The deposited data will be publicly available as of the publication date. Custom R scripts used for data analysis in this study have been developed in our lab and are available upon request.

## ACKNOWLEDGEMENTS

We are grateful to Paul Zumbo, Friederike Dundar, and the Applied Bioinformatics Core at Weill Cornell Medicine for establishing STARmulti analysis, William Lai (Cornell Ithaca) for advice on ChIP-exo, Matthew Lorincz and Julie Brind’Amour for advice on ULI-NChIP-seq, Charlene Raclot and Didier Trono for sharing DPPA2 and DPPA4 KO mESCs and Wee Wei Tee for sharing gene expression data. We thank all members of the Apostolou and Stadtfeld labs for their valuable comments and support. C.M.U. was supported by fellowships from the New York State Department of Health (NYSTEM training program) C32558GG and the National Heart, Lung, and Blood Institute (NHLBI) T32HL160520. M.S. was supported by grants from the NIH (R01GM145864), the Simons Foundation, the Tri-Institutional Stem Cell Initiative (Tri-SCI), and the Bohmfalk Charitable Trust. E.A. was funded by NIH (5R01GM138635 and 5RM1GM139738).

## AUTHOR CONTRIBUTIONS

K.C. established and validated cell lines, conducted and analyzed cell sorting, immunofluorescence, and WB experiments, and prepared cells and material for genomics assays. C.M.U. conducted all bioinformatics analyses and assisted with CUT&RUN experiments. K.K. conducted H3K9me2 ULI-NChIP-seq and intracellular flow cytometry experiments. S.M. assisted in characterizing cell lines and in ChIP-exo experiments. L.S. conducted ATAC-seq experiments. A.P. advised on initial bioinformatic analyses. M.S. assisted in gene targeting experiments. The study was initially conceived by K.C. and M.S. and further conceptualized with C.M.U. and E.A. All experiments were planned by K.C. and C.M.U. with advice and supervision from M.S. and E.A. The manuscript was written by M.S., C.M.U., and K.C. and edited by K.K. and E.A., with input from all authors. M.S. and E.A. acquired funding.

## SUPPLEMENTAL INFORMATION TITLES AND LEGENDS

**TABLE S1:** RNA-Seq gene tables for all datasets.

**TABLE S2:** List of ECORDs.

**TABLE S3:** ChromHMM reference tables.

**TABLE S4:** Chromatin Binding information for EHMT2 and DPPA4.

**TABLE S5:** Gene ontology terms and genes.

**TABLE S6:** Supplementary materials.

**TABLE S7:** Statistical Analyses.

## REFERENCES

1. Baedke, J. The epigenetic landscape in the course of time: Conrad Hal Waddington’s methodological impact on the life sciences. Stud Hist Philos Biol Biomed Sci 44, 756–73 (2013).

2. Macfarlan, T.S. et al. Embryonic stem cell potency fluctuates with endogenous retrovirus activity. Nature 487, 57–63 (2012).

3. De Iaco, A., Coudray, A., Duc, J. & Trono, D. DPPA2 and DPPA4 are necessary to establish a 2C-like state in mouse embryonic stem cells. EMBO Rep 20(2019).

4. Eckersley-Maslin, M. et al. Dppa2 and Dppa4 directly regulate the Dux-driven zygotic transcriptional program. Genes Dev 33, 194–208 (2019).

5. Hendrickson, P.G. et al. Conserved roles of mouse DUX and human DUX4 in activating cleavage-stage genes and MERVL/HERVL retrotransposons. Nat Genet 49, 925–934 (2017).

6. Yang, F. et al. DUX-miR-344-ZMYM2-Mediated Activation of MERVL LTRs Induces a Totipotent 2C-like State. Cell Stem Cell 26, 234–250 e7 (2020).

7. Jerabek, S., Merino, F., Scholer, H.R. & Cojocaru, V. OCT4: dynamic DNA binding pioneers stem cell pluripotency. Biochim Biophys Acta 1839, 138–54 (2014).

8. Nakatani, T. & Torres-Padilla, M.E. Regulation of mammalian totipotency: a molecular perspective from in vivo and in vitro studies. Curr Opin Genet Dev 81, 102083 (2023).

9. Jia, Y.K., Yu, Y. & Guan, L. Advances in understanding the regulation of pluripotency fate transition in embryonic stem cells. Front Cell Dev Biol 12, 1494398 (2024).

10. Boskovic, A. et al. Higher chromatin mobility supports totipotency and precedes pluripotency in vivo. Genes Dev 28, 1042–7 (2014).

11. Eckersley-Maslin, M.A. et al. MERVL/Zscan4 Network Activation Results in Transient Genome-wide DNA Demethylation of mESCs. Cell Rep 17, 179–192 (2016).

12. Zhu, Y. et al. Relaxed 3D genome conformation facilitates the pluripotent to totipotent-like state transition in embryonic stem cells. Nucleic Acids Res 49, 12167–12177 (2021).

13. Tachibana, M., Sugimoto, K., Fukushima, T. & Shinkai, Y. Set domain-containing protein, G9a, is a novel lysine-preferring mammalian histone methyltransferase with hyperactivity and specific selectivity to lysines 9 and 27 of histone H3. J Biol Chem 276, 25309–17 (2001).

14. Rice, J.C. et al. Histone methyltransferases direct different degrees of methylation to define distinct chromatin domains. Mol Cell 12, 1591–8 (2003).

15. Tachibana, M. et al. Histone methyltransferases G9a and GLP form heteromeric complexes and are both crucial for methylation of euchromatin at H3-K9. Genes Dev 19, 815–26 (2005).

16. Tachibana, M. et al. G9a histone methyltransferase plays a dominant role in euchromatic histone H3 lysine 9 methylation and is essential for early embryogenesis. Genes Dev 16, 1779–91 (2002).

17. Zylicz, J.J. et al. G9a regulates temporal preimplantation developmental program and lineage segregation in blastocyst. Elife 7(2018).

18. Yelagandula, R. et al. ZFP462 safeguards neural lineage specification by targeting G9A/GLP- mediated heterochromatin to silence enhancers. Nat Cell Biol 25, 42–55 (2023).

19. Mozzetta, C. et al. The histone H3 lysine 9 methyltransferases G9a and GLP regulate polycomb repressive complex 2-mediated gene silencing. Mol Cell 53, 277–89 (2014).

20. Maksakova, I.A. et al. Distinct roles of KAP1, HP1 and G9a/GLP in silencing of the two-cell-specific retrotransposon MERVL in mouse ES cells. Epigenetics Chromatin 6, 15 (2013).

21. Nabet, B. et al. The dTAG system for immediate and target-specific protein degradation. Nat Chem Biol 14, 431–441 (2018).

22. Shinkai, Y. & Tachibana, M. H3K9 methyltransferase G9a and the related molecule GLP. Genes Dev 25, 781–8 (2011).

23. Tachibana, M., Matsumura, Y., Fukuda, M., Kimura, H. & Shinkai, Y. G9a/GLP complexes independently mediate H3K9 and DNA methylation to silence transcription. EMBO J 27, 2681–90 (2008).

24. Di Giammartino, D.C. et al. KLF4 is involved in the organization and regulation of pluripotency-associated three-dimensional enhancer networks. Nat Cell Biol 21, 1179–1190 (2019).

25. Ernst, J. & Kellis, M. Chromatin-state discovery and genome annotation with ChromHMM. Nat Protoc 12, 2478–2492 (2017).

26. Vu, H. & Ernst, J. Universal chromatin state annotation of the mouse genome. Genome Biol 24, 153 (2023).

27. Lieberman-Aiden, E. et al. Comprehensive mapping of long-range interactions reveals folding principles of the human genome. Science 326, 289–93 (2009).

28. Falco, G. et al. Zscan4: a novel gene expressed exclusively in late 2-cell embryos and embryonic stem cells. Dev Biol 307, 539–50 (2007).

29. Ji, S. et al. OBOX regulates mouse zygotic genome activation and early development. Nature 620, 1047–1053 (2023).

30. Wu, J. et al. The landscape of accessible chromatin in mammalian preimplantation embryos. Nature 534, 652–7 (2016).

31. Ishiuchi, T. et al. Early embryonic-like cells are induced by downregulating replication-dependent chromatin assembly. Nat Struct Mol Biol 22, 662–71 (2015).

32. Polo, J.M. et al. A molecular roadmap of reprogramming somatic cells into iPS cells. Cell 151, 1617–32 (2012).

33. Shen, H. et al. Mouse totipotent stem cells captured and maintained through spliceosomal repression. Cell 184, 2843–2859 e20 (2021).

34. Iturbide, A. et al. Retinoic acid signaling is critical during the totipotency window in early mammalian development. Nat Struct Mol Biol 28, 521–532 (2021).

35. Rossi, M.J., Lai, W.K.M. & Pugh, B.F. Simplified ChIP-exo assays. Nat Commun 9, 2842 (2018).

36. Skene, P.J. & Henikoff, S. An efficient targeted nuclease strategy for high-resolution mapping of DNA binding sites. Elife 6(2017).

37. Teissandier, A., Servant, N., Barillot, E. & Bourc’his, D. Tools and best practices for retrotransposon analysis using high-throughput sequencing data. Mob DNA 10, 52 (2019).

38. Brind’Amour, J. et al. An ultra-low-input native ChIP-seq protocol for genome-wide profiling of rare cell populations. Nat Commun 6, 6033 (2015).

39. Brind’Amour, J. & Lorincz, M.C. Profiling Histone Methylation in Low Numbers of Cells. Methods Mol Biol 2529, 229–251 (2022).

40. Rowe, H.M. et al. KAP1 controls endogenous retroviruses in embryonic stem cells. Nature 463, 237–40 (2010).

41. Bian, C., Chen, Q. & Yu, X. The zinc finger proteins ZNF644 and WIZ regulate the G9a/GLP complex for gene repression. Elife 4(2015).

42. Yan, Z. et al. G9a/GLP-sensitivity of H3K9me2 Demarcates Two Types of Genomic Compartments. Genomics Proteomics Bioinformatics 18, 359–370 (2020).

43. Fukuda, K. et al. Regulation of mammalian 3D genome organization and histone H3K9 dimethylation by H3K9 methyltransferases. Commun Biol 4, 571 (2021).

44. Zhang, W. et al. Zscan4c activates endogenous retrovirus MERVL and cleavage embryo genes. Nucleic Acids Res 47, 8485–8501 (2019).

45. Fort, A. et al. Deep transcriptome profiling of mammalian stem cells supports a regulatory role for retrotransposons in pluripotency maintenance. Nat Genet 46, 558–66 (2014).

46. Peters, A.H. et al. Partitioning and plasticity of repressive histone methylation states in mammalian chromatin. Mol Cell 12, 1577–89 (2003).

47. Spracklin, G. et al. Diverse silent chromatin states modulate genome compartmentalization and loop extrusion barriers. Nat Struct Mol Biol 30, 38–51 (2023).

48. Vega-Sendino, M. et al. The homeobox transcription factor DUXBL controls exit from totipotency. Nat Genet 56, 697–709 (2024).

49. Hubert, K.A. & Wellik, D.M. Hox genes in development and beyond. Development 150(2023).

50. Jachowicz, J.W. et al. LINE-1 activation after fertilization regulates global chromatin accessibility in the early mouse embryo. Nat Genet 49, 1502–1510 (2017).

51. Li, X. et al. LINE-1 transcription activates long-range gene expression. Nat Genet 56, 1494–1502 (2024).

52. Kleefstra, T. & de Leeuw, N. Kleefstra Syndrome. in GeneReviews((R)) (eds. Adam, M.P. et al.) (Seattle (WA), 1993).

53. Roopra, A., Qazi, R., Schoenike, B., Daley, T.J. & Morrison, J.F. Localized domains of G9a-mediated histone methylation are required for silencing of neuronal genes. Mol Cell 14, 727–38 (2004).

54. Beard, C., Hochedlinger, K., Plath, K., Wutz, A. & Jaenisch, R. Efficient method to generate single-copy transgenic mice by site-specific integration in embryonic stem cells. Genesis 44, 23–8 (2006).

55. Schneider, C.A., Rasband, W.S. & Eliceiri, K.W. NIH Image to ImageJ: 25 years of image analysis. Nat Methods 9, 671–5 (2012).

56. Kassambara. rstatix: Pipe-Friendly Framework for Basic Statistical Tests. R package version 0.7.2, https://rpkgs.datanovia.com/rstatix/. (2023).

57. Buenrostro, J.D., Wu, B., Chang, H.Y. & Greenleaf, W.J. ATAC-seq: A Method for Assaying Chromatin Accessibility Genome-Wide. Curr Protoc Mol Biol 109, 21 29 1–21 29 9 (2015).

58. Pelham-Webb, B. et al. H3K27ac bookmarking promotes rapid post-mitotic activation of the pluripotent stem cell program without impacting 3D chromatin reorganization. Mol Cell 81, 1732–1748 e8 (2021).

59. Skene, P.J., Henikoff, J.G. & Henikoff, S. Targeted in situ genome-wide profiling with high efficiency for low cell numbers. Nat Protoc 13, 1006–1019 (2018).

60. Ee, L.-s., et al. Transcriptional remodeling by OTX2 directs specification and patterning of mammalian definitive endoderm. bioRxiv (2024).

61. FelixKrueger/TrimGalore: A wrapper around Cutadapt and FastQC to consistently apply adapter and quality trimming to FastQ files, with extra functionality for RRBS data.

62. Krueger, F. TrimGalore: A wrapper around Cutadapt and FastQC to consistently apply adapter and quality trimming to FastQ files, with extra functionality for RRBS data.

63. Dobin, A. et al. STAR: ultrafast universal RNA-seq aligner. Bioinformatics 29, 15–21 (2013).

64. Jin, Y., Tam, O.H., Paniagua, E. & Hammell, M. TEtranscripts: a package for including transposable elements in differential expression analysis of RNA-seq datasets. Bioinformatics 31, 3593–9 (2015).

65. Frankish, A. et al. GENCODE reference annotation for the human and mouse genomes. Nucleic Acids Res 47, D766–D773 (2019).

66. Ramirez, F., Dundar, F., Diehl, S., Gruning, B.A. & Manke, T. deepTools: a flexible platform for exploring deep-sequencing data. Nucleic Acids Res 42, W187–91 (2014).

67. Love, M.I., Huber, W. & Anders, S. Moderated estimation of fold change and dispersion for RNA-seq data with DESeq2. Genome Biol 15, 550 (2014).

68. Budiaji, W. kmed: Distance-Based k-Medoids. (2022).

69. Hu, Z. et al. Maternal factor NELFA drives a 2C-like state in mouse embryonic stem cells. Nat Cell Biol 22, 175–186 (2020).

70. Wickham, H. A Layered Grammar of Graphics. Journal of Computational and Graphical Statistics 19, 3–28 (2010).

71. Ge, S.X., Jung, D. & Yao, R. ShinyGO: a graphical gene-set enrichment tool for animals and plants. Bioinformatics 36, 2628–2629 (2020).

72. Sayols, S. rrvgo: a Bioconductor package for interpreting lists of Gene Ontology terms. MicroPubl Biol 2023(2023).

73. Tools, P. Picard Tools - By Broad Institute.

74. Li, H. et al. The Sequence Alignment/Map format and SAMtools. Bioinformatics (Oxford, England) 25, 2078–2079 (2009).

75. Quinlan, A.R. & Hall, I.M. BEDTools: a flexible suite of utilities for comparing genomic features. Bioinformatics 26, 841–2 (2010).

76. Zhang, Y. et al. Model-based analysis of ChIP-Seq (MACS). Genome Biol 9, R137 (2008).

77. Au Yeung, W.K., et al. Histone H3K9 Methyltransferase G9a in Oocytes Is Essential for Preimplantation Development but Dispensable for CG Methylation Protection. Cell reports 27, 282–293.e4 (2019).

78. Lawrence, M. et al. Software for computing and annotating genomic ranges. PLoS computational biology 9, e1003118 (2013).

79. Lee, S., Cook, D. & Lawrence, M. plyranges: a grammar of genomic data transformation. Genome biology 20, 4 (2019).

80. Lawrence, M., Gentleman, R. & Carey, V. rtracklayer: an R package for interfacing with genome browsers. Bioinformatics (Oxford, England) 25, 1841–1842 (2009).

81. Carlson, M. org.Mm.eg.db: Genome wide annotation for Mouse, (2021).

82. Jin, Y., Tam, O.H., Paniagua, E. & Hammell, M. TEtranscripts: a package for including transposable elements in differential expression analysis of RNA-seq datasets. Bioinformatics (Oxford, England) 31, 3593–3599 (2015).

83. Stempor, P. & Ahringer, J. SeqPlots - Interactive software for exploratory data analyses, pattern discovery and visualization in genomics. Wellcome open research 1, 14 (2016).

84. BBTools. BBTools. DOE Joint Genome Institute.

85. Buenrostro, J.D., Wu, B., Chang, H.Y. & Greenleaf, W.J. ATAC-seq: A Method for Assaying Chromatin Accessibility Genome-Wide. Current protocols in molecular biology 109, 21.29.1–21.29.9 (2015).

86. Mölder, F. et al. Sustainable data analysis with Snakemake. F1000Research 10, 33 (2021).

87. Dobin, A. et al. STAR: ultrafast universal RNA-seq aligner. Bioinformatics (Oxford, England) 29, 15–21 (2013).

88. Zhu, L.J. et al. ChIPpeakAnno: a Bioconductor package to annotate ChIP-seq and ChIP-chip data. BMC bioinformatics 11, 237 (2010).

89. Wickham, H. et al. Welcome to the Tidyverse. Journal of Open Source Software 4, 1686 (2019).

90. Pedersen, T.L. patchwork: The Composer of Plots, (2024).

91. Larsson, J. eulerr: Area-Proportional Euler and Venn Diagrams with Ellipses, (2024).

92. Krassowski, M. ComplexUpset. (2020).

93. Schiettekatte, N., Brandl, S. & Casey, J. fishualize: Color Palettes Based on Fish Species, (2019).

94. Brunson, J.C. & Read, Q.D. ggalluvial: Alluvial Plots in ‘ggplot2’.

