## Supplemental Figures and Legends for "Coordinated repression of totipotency-associated gene loci by histone methyltransferase EHMT2 through binding to LINE-1 regulatory elements"

**Lead contact**

FIGURE S1

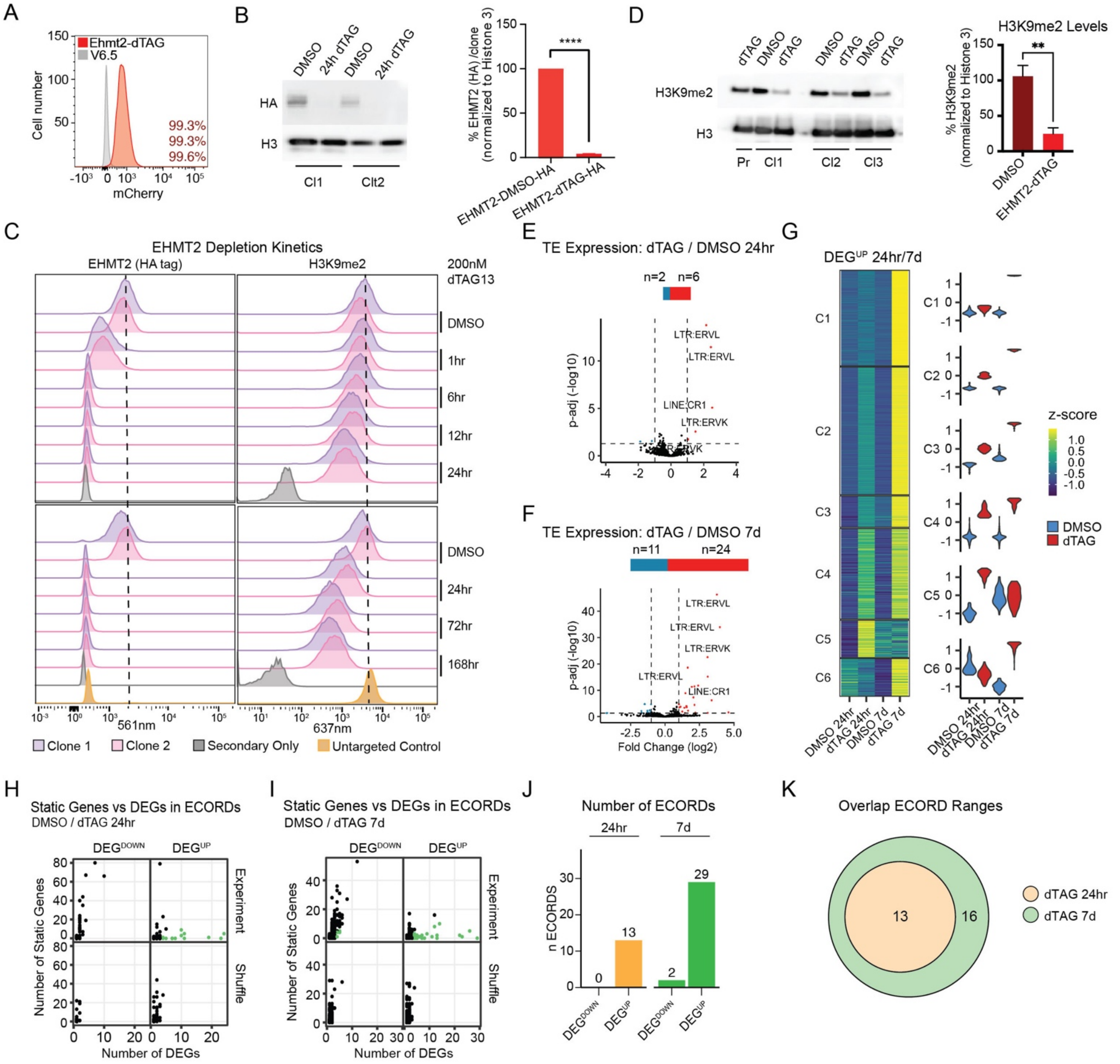


**Figure S1. Acute EHMT2 depletion reveals distinct categories of target genes. Related to Figure 1.**

**A)** Flow cytometry of EHMT2 dTAG lines. mCherry acts as a transcriptional reporter. Percentages indicate expression levels of 3 selected clones. **B)** Western blot of EHMT2 (HA) levels after 24h dTAG treatment. ****p<0.0001; two-tailed, paired t-test. Error bars represent mean with SD. **C)** Flow cytometry time course of EHMT2 and H3K9me2 levels during continuous dTAG treatment. **D)** Western blot of H3K9me2 levels after 24h dTAG treatment. **p<0.01; two-tailed, unpaired t-test. Error bars represent mean with SD. **E-F)** Volcano plots showing numbers of differentially expressed (DE) repetitive elements in response to 24h and 7d continuous dTAG treatment, respectively. DE: Adj. p-value <0.05 & absolute log2 fold change ≥1 **G)** K-medoids clustering of all DE genes after 24h or 7d dTAG. Genes are z-scored by row. **H-I)** Number of static genes in ECORDs at 24h and 7d, respectively. Green indicates ECORDS ≥4 DEGs. **J)** The number of ECORDs at 24h and 7d. **K)** Overlap of ECORD ranges. For ECORD ranges in H-I, the furthest start and end coordinate of all genes in the ECORD was used. Error bars indicate mean.

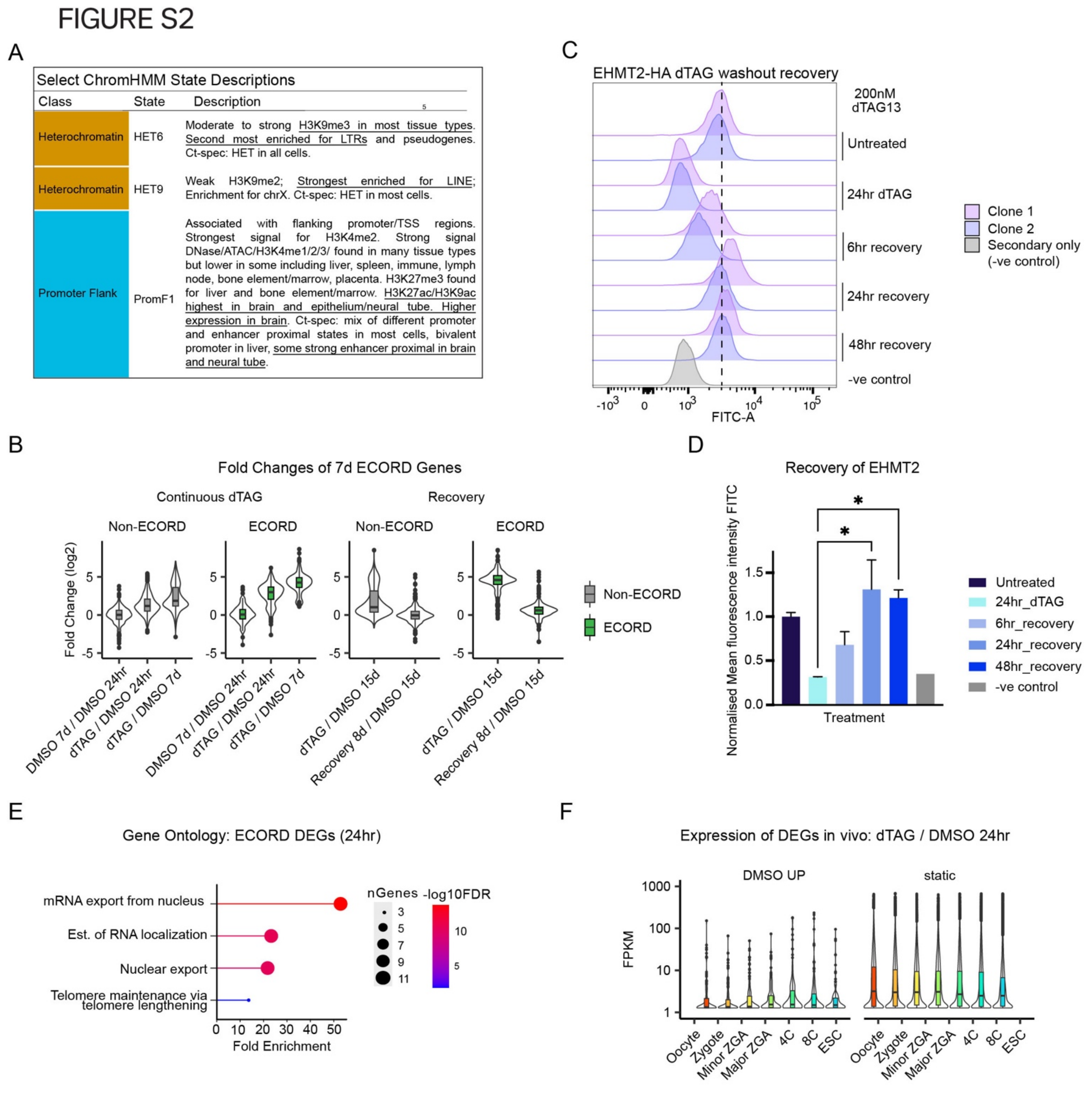


**Figure S2. Figure 2. Genes in ECORDs have distinct properties and are enriched for genes involved in Zygotic Genome Activation (ZGA). Related to Figure 2.**

**A)** The descriptions of ChromHMM terms shown in Fig.2A. **B)** Fold-changes of 7d dTAG Up DEGs after 15d continuous dTAG treatment (left) and after 7d continuous dTAG treatment followed by 8d of dTAG washout ("Recovery", right). **C)** Flow cytometry time course showing EHMT2 (HA) recovery after dTAG washout. A sample treated with secondary antibody only was used as a negative (-ve) control. **D)** Quantification of mean fluorescence intensity from C.

*p<0.05; One way ANOVA. Error bars represent mean with SD. **E)** Gene ontology (GO) terms of genes up-regulated after 24h dTAG treatment for ECORD DEGs. **F)** Expression of 24h DMSO Up DEGs and static genes in an *in vivo* time course of embryonic development (Wu et al., 2016). Only genes with an FPKM ≥10 in at least one sample type were included.

FIGURE S3

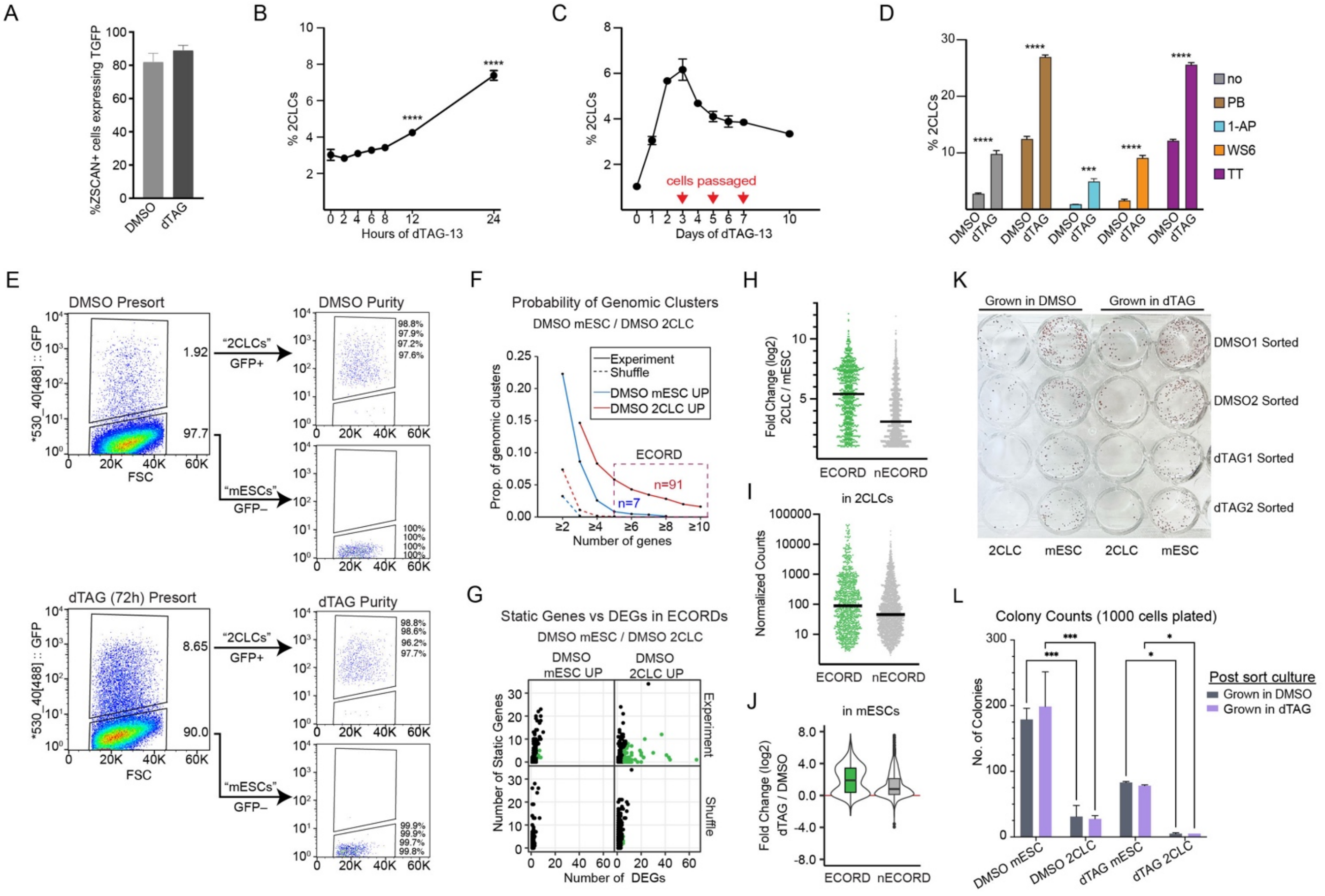


**Figure S3. EHMT2 acts as a gatekeeper for entry into the 2-Cell Like State (2CLC). Related to Figure 3.**

**A)** Quantification of the percentage of ZSCAN4 positive cells that express turboGFP in DMSO and dTAG conditions. Error bars represent the mean with SD. **B)** Percentage of 2CLCs in mESC cultures after indicated hours of dTAG-13 treatment compared to DMSO treated (0h) cells. ****p<0.0001 with one-way ANOVA. Shown is mean with SD. **C)** Percentage of 2CLCs after indicated days of dTAG-13 treatment. Shown is mean with SD. Red arrows indicate when mESC cultures were passaged. **D)** Percentage of 2CLCs after 24h of mESC culture in the presence of the indicated chemicals. Shown is mean with SD. Nothing added, no; Pladienolide B, PB (spliceosome inhibitor); 1-azakenpaullone, 1-AP (GSK3 inhibitor); WS6 (IkB kinase and EPB1 inhibitor); TTNPB, TT (RAR agonist). ***p<0.001, ****p<0.0001; multiple t-tests using Holm-Sidak correction. **E)** FACS analysis indicating percentage 2CLC and mESC presorting after treatment with DMSO or dTAG. Post-sorting purity and gates are indicated for each gated population. **F)** Probability curves showing the fraction of gene clusters with ≥n DEGs and < 50%

static genes that do not break TAD boundaries. For sorted RNAseq experiments, “ECORDs” were defined as clusters with n ≥5 DEGs and <0.5 proportion of static genes. **G)** Number of static genes in ECORDs at 24h and 7d, respectively. Green indicates ECORDS ≥5 DEGs. **H)** Fold change of ECORD and Non-ECORD Up DEGs (DMSO 2CLC > mESC). **I)** Normalized expression of ECORD and Non-ECORD Up DEGS (DMSO 2CLC > mESC) in DMSO 2CLC **J)** Fold changes of ECORD and Non-ECORD Up DEGs (mESC dTAG/DMSO) in mESC dTAG (mESC dTAG / DMSO). **K)** Plate showing Alkaline phosphatase-stained colonies after 6 days of plating of sorted populations in different media conditions. **L)** Quantification of the colonies shown in panel K. *p<0.05, ***p<0.001; two-way ANOVA. Error bars represent the mean with SD.

FIGURE S4

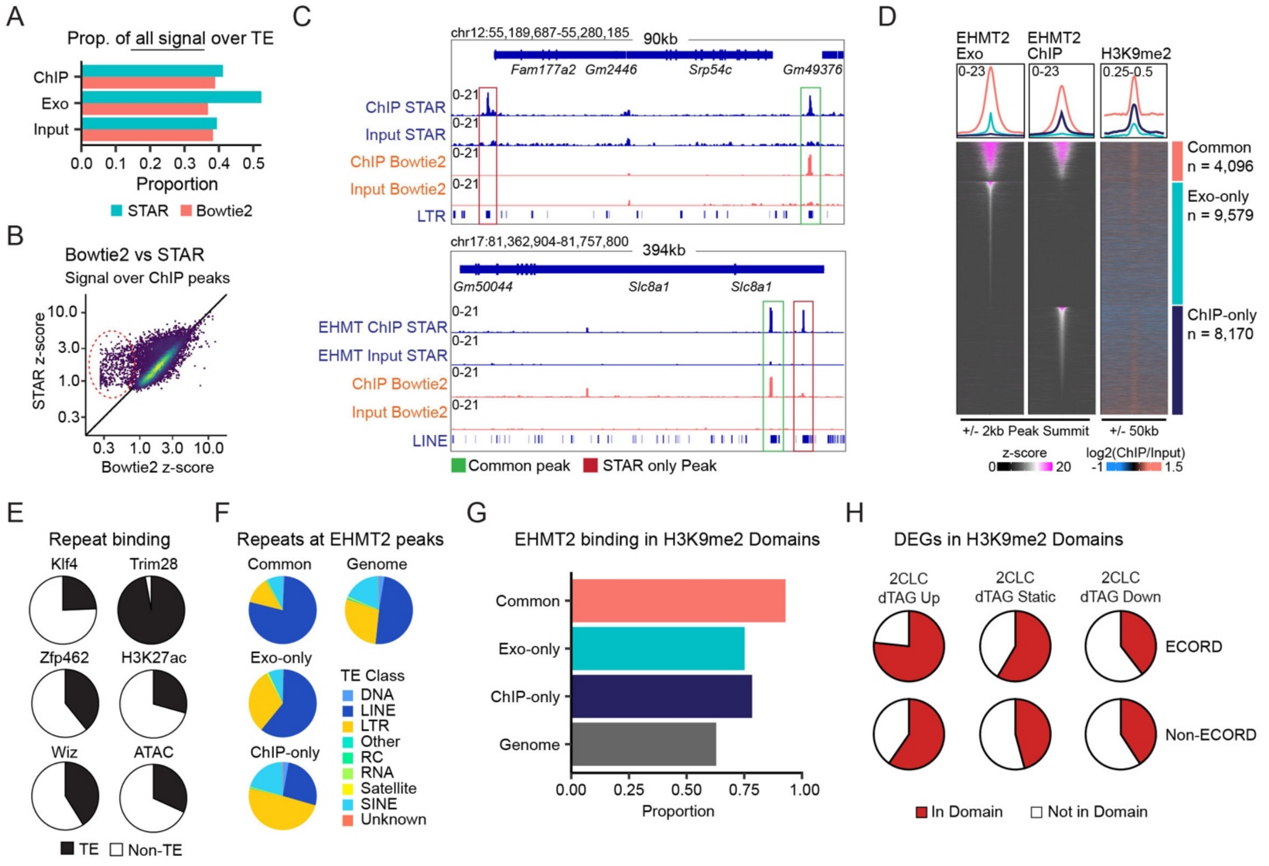


**Figure S4. H3K9me2 domains repress ECORD expression. Related to Figure 4.**

**A)** Comparison of Bowtie2 and modified STAR multi-alignment (see methods) showing the proportion of all signal over repeats. **B)** Correlation of EHMT2 ChIP signal after aligning with Bowtie2 or STARmulti (see methods). Most peaks are unaffected, but a subset of *de novo* peaks can be identified with STARmulti (dashed circle). **C)** Examples of peaks only identified using STARmulti (red) and peaks unaffected by the aligner used (green). **D)** EHMT2 and H3K9me2 signal around summits of ChIP-Exo and ChIP peaks. For common peaks, the mid-point of the summit was used. **E)** Proportion of peaks that overlap repetitive elements. Proportion was calculated at the base pair level. **F)** Classes of repetitive elements (TEs) that EHMT2 binds. **G)** Proportion of EHMT2 ChIP and ChIP-Exo peaks that overlap an H3K9me2 domain. **H)** Proportion of RNAseq k-meds genes that overlap an H3K9me2 domain split by whether they fall into an ECORD called in DMSO 2CLC / mESC.

FIGURE S5

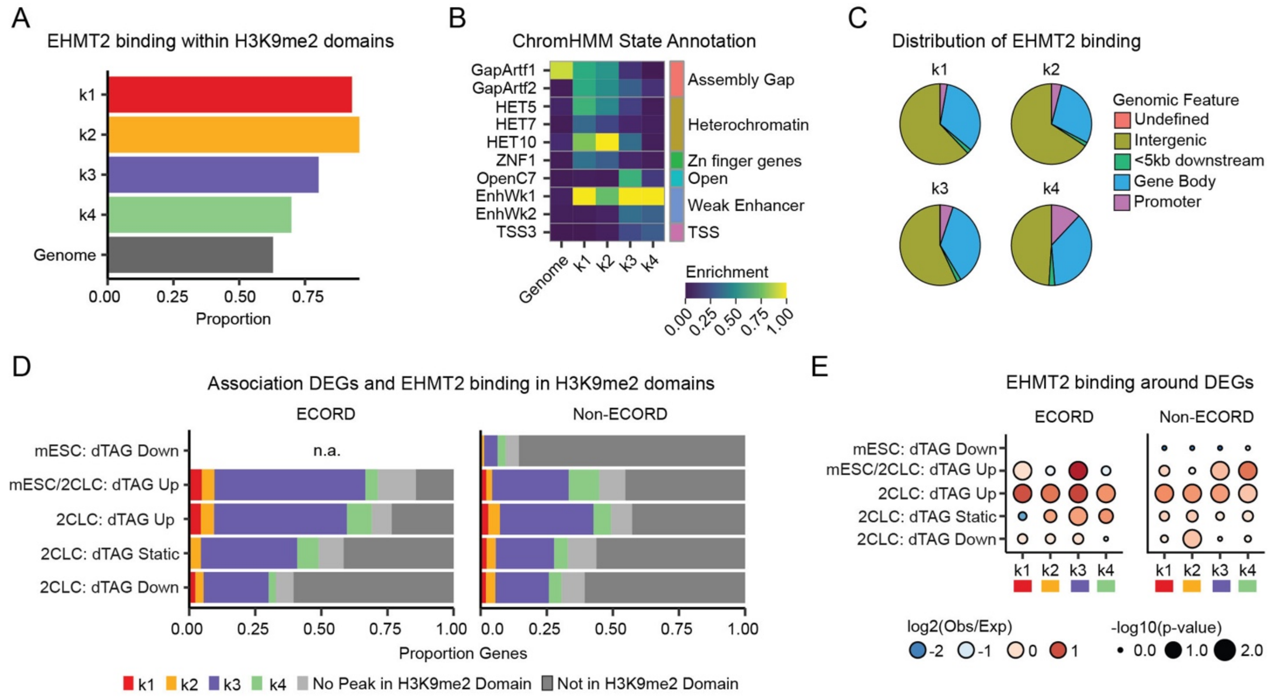


**Figure S5. EHMT2 binding suggests locus-specific modes of gene regulation. Related to Figure 5.**

**A)** Proportion of peaks in EHMT2 k-means clusters that overlap H3K9me2 domains. **B)** ChromHMM of EHMT2 peaks using a 100-state model (Vu and Ernst, 2023). **C)** Genomic distribution of peaks in EHMT2 k-means clusters. **D)** Enrichment of EHMT2 k-means clusters with DEGs. DEGs were assigned to the nearest EHMT2 peak in the same H3K9me2 domain. P-value was calculated using a bootstrap approach by shuffling gene categories 1000 times. **E)** Association between RNA k-meds clusters, ECORDs, and EHMT2 binding. Genes were assigned to the nearest EHMT2 peak in the same H3K9me2 domain.

FIGURE S6

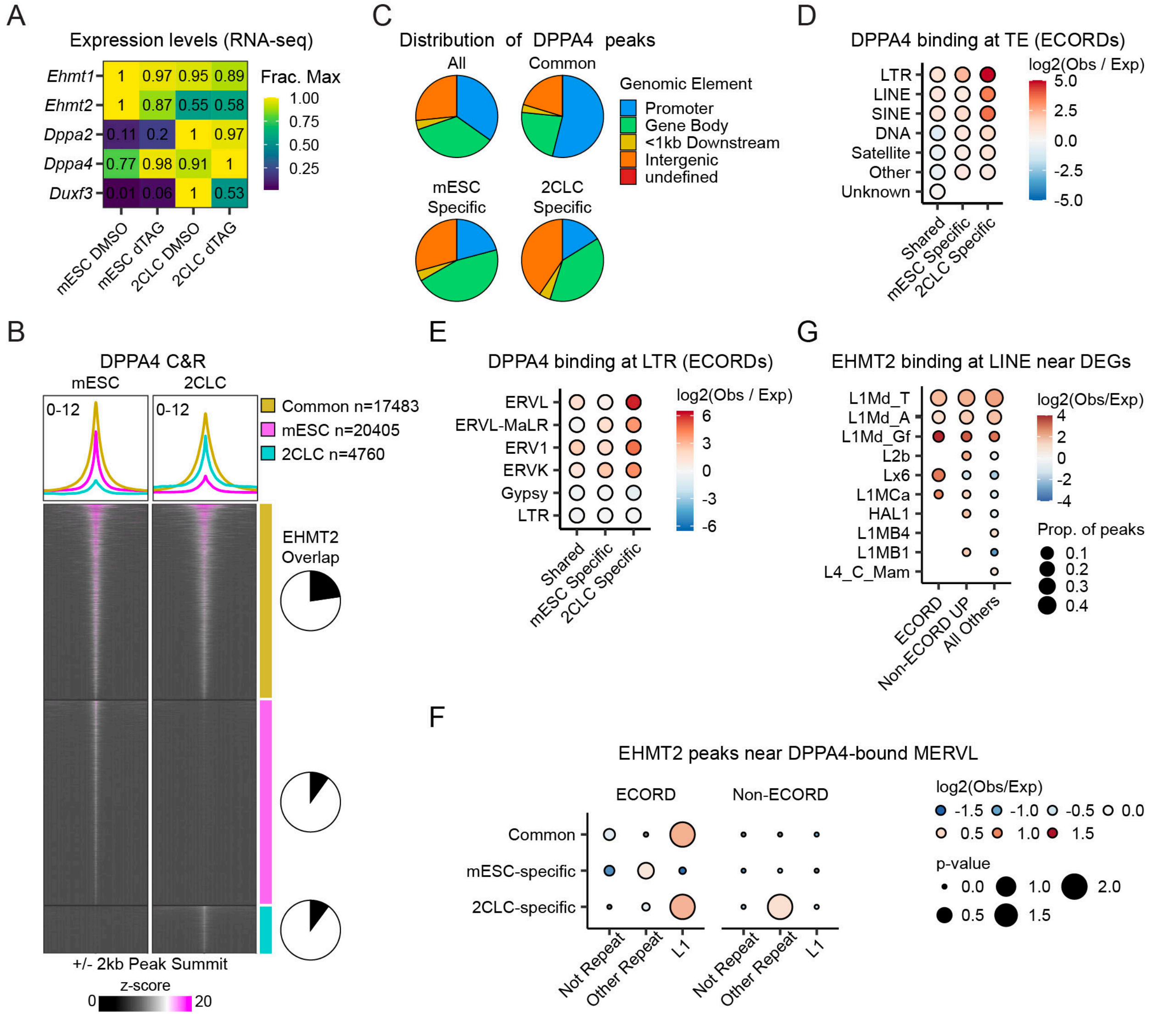


**Figure S6. EHMT2 antagonizes DPPA2/4 function at ECORDs. Related to Figure 6.**

**A)** Expression of *Ehmt1/2*, *Dppa2/4,* and *Dux* in sorted RNAseq. **B)** DPPA4 C&R signal (+/-2kb of peak summits) around all DPPA4 peaks. For common peaks, the mid-point of the summits was used. **C)** Genomic distribution of DPPA4 peaks. **D)** Observed vs expected ratio (log2) of DPPA4 peaks over different classes of repetitive elements (TEs). **E)** Observed vs expected ratio (log2) of DPPA4 peaks over different families of LTRs. For this analysis, only peaks that overlapped an LTR were used. The expected ratio was calculated for D and E by shuffling Dppa4 peaks over the genome 100 times. **F)** Overlap of all DPPA4 and DUX peaks. **G)** Observed vs expected ratio (log2) of LINE repeat genes that EHMT2 binds to split by the type of DEG they are associated with. To associate peaks and DEGs, each DEG was assigned to the nearest

EHMT2 peak in the same H3K9me2 domain (see methods). **I)** Enrichment of EHMT2 peaks around Dppa4 MERVL at ECORDs and Non-ECORD UP DEGs. Dppa4 peaks were assigned to the nearest EHMT2 peak in the same H3K9me2 domain. P-value was calculated using a bootstrap approach by shuffling Dppa4 peaks over the genome 1000 times.
